# DRG meningeal tertiary lymphoid structures are regulated by B cells as a pronociceptive locus after peripheral nerve injury

**DOI:** 10.64898/2026.08.09.743585

**Authors:** Tusar K. Acharya, Vipul K. Pandey, Kendal F. Willcox, Nathan T. Fiore, Gabriel V. Lucena-Silva, Jayden A. O’Brien, Allison M. Barry, Joseph B. Lesnak, Sever M. Zagrai, David M. Ruiz, Younus A. Zuberi, Michael J. Lacagnina, Pooja Singhmar, Lisa M.F. Janssen, Abbie V. Viscardi, Rachel E. Miller, Anne-Marie Malfait, Martin K. Lotz, Rajasekaran Mahalingam, Hans F. Coetzee, Theodore J. Price, Thiago M. Cunha, Cobi J. Heijnen, Peter M. Grace

**Affiliations:** Department of Translational Neuroscience, UT MD Anderson, Houston, Texas 77030, USA; Center for Research in Inflammatory Diseases (CRID), Ribeirão Preto Medical School, University of Sao Paulo, Ribeirão Preto, Sao Paulo 14040-900, Brazil; Department of Neuroscience and Center for Advanced Pain Studies, University of Texas at Dallas, Richardson, Texas, 75080, USA; Department of Anesthesia, Cincinnati Children’s Hospital Medical Center, Cincinnati, Ohio, 45229, USA; Department of Molecular Medicine, Scripps Research, La Jolla, California, 92037, USA; Department of Anatomy and Physiology, College of Veterinary Medicine, Kansas State University, Manhattan, Kansas, 66506, USA; Department of Internal Medicine, Division of Rheumatology, Rush University Medical Center, Chicago, Illinois, 60612, USA; IMmune HEAlth International Research Laboratory (IMHEA-IRL), French National Centre for Scientific Research (CNRS) at Ribeirão Preto Medical School, University of Sao Paulo, Brazil; Department of Psychological Sciences, Rice University, Houston, Texas, 77005, USA; Cancer Neuroscience Program, UT MD Anderson, Houston, Texas 77030, USA

**Keywords:** Ectopic lymphoid structures, tertiary lymphoid organs, meninges, neuropathy, neuropathic pain

## Abstract

B cell-derived IgG in the dorsal root ganglia (DRG) drives neuropathic pain after peripheral nerve injury (PNI), but the site of B cell organization is unclear. Here, PNI induced leukocyte clusters in the DRG meninges, enveloped by lymphatic endothelium and apposed to high endothelial venules. These clusters resemble tertiary lymphoid structures (TLSs) with germinal center-like features, including germinal center B cells and plasma cells, and follicular dendritic and follicular helper T cells. Single-cell RNA sequencing revealed enrichment of germinal center B cells in the DRG meninges after PNI. Germinal center B cells regulate TLS organization: TLSs were absent after deletion of *Ezh2* from germinal center-experienced B cells. Intrathecal CD20 monoclonal antibody to locally deplete B cells also disrupted TLS organization. Conversely, intrathecal B cell transfer to B cell-deficient (muMT) mice was sufficient for TLS organization after PNI. Allodynia did not develop when TLS organization was disordered. Similar TLSs formed in pig DRG after tail docking and in human donors with chronic pain, where B cell receptor clonotype analysis confirmed functional maturity. Together, these data establish that germinal center B cells are required for TLS organization, and that disrupting this process abolishes the development of neuropathic pain after PNI.

## INTRODUCTION

Peripheral nerve injury (PNI) provokes a coordinated innate and adaptive immune response which contributes to neuropathic pain, a debilitating and intractable neurological condition^1, 2, 3, 4, 5, 6^. New evidence suggests that immune cells in the meninges of the dorsal root ganglia (DRG) are unique contributors to neuropathic pain. Dendritic cells and CD4^+^ T helper cells accumulate at this site after PNI and secrete peptides and other factors to excite spinal cord nociceptive neurons^7, 8^. T helper cells activate B cells, which we recently implicated in PNI-induced pain through a mechanism involving immunoglobulin G (IgG) secretion^9, 10, 11^. IgG accumulates throughout the DRG parenchyma where it activates Fc gamma receptors to drive neuronal hyperexcitability and pain^9, 10, 11^. Importantly, IgG accumulation is conserved in humans, with elevated deposition observed in DRG samples from donors with chronic pain^9^. Although single-cell RNA sequencing identified a population of B cells in mouse and human DRG, as well as enrichment of a B cell transcriptional signature in DRGs from human donors with neuropathic pain, their numbers in the parenchyma were low and were not altered following PNI^9^. Given evidence of antigen-presenting cells and T cell accumulation in the meninges of the DRG after PNI^7, 8, 12^, it is possible that B cells may also be activated and differentiated at the meningeal site.

It is well known that the final stages of B cell differentiation into antibody-secreting cells can take place in so-called germinal centers of the secondary lymphoid organs. Here, (auto)antigen-activated B cells can interact with their cognate CD4 follicular T helper (Tfh) cells. As a result, B cells undergo clonal expansion and somatic hypermutation of their immunoglobulin loci, before terminally differentiating into plasma cells or memory B cells^13, 14^. In addition, terminal differentiation of B cells can take place in organized lymphoid aggregates termed tertiary lymphoid structures (TLSs) that are reminiscent of germinal centers and develop in the perivascular space in response to disturbed tissue homeostasis^15, 16, 17, 18^.

In patients with autoimmune diseases or other chronic inflammatory conditions, the presence of TLSs is often associated with more severe pathology^17, 18, 19^. TLSs in the spinal cord meninges have been observed in preclinical models of spinal pathology, including neurodegenerative disease (amyotrophic lateral sclerosis, experimental autoimmune encephalomyelitis) and trauma (spinal cord injury)^20^. Whether TLSs are formed in the peripheral nervous system and contribute to neuropathic pain remains to be determined. Here, we identify TLSs in the DRG meninges following PNI, characterize their cellular and molecular composition, and demonstrate that they are conserved in pigs and humans. Using complementary approaches—local B cell depletion, adoptive transfer, conditional genetic deletion, single-cell and spatial transcriptomics—we establish that germinal center B cells are required for TLS organization and that disrupting this process abolishes the development of neuropathic pain after PNI.

## RESULTS

### PNI induces TLSs in the meninges of the DRG

We investigated whether TLSs would emerge in the meninges of the ipsilateral L4/5 DRG because of sciatic PNI (chronic constriction injury). We performed whole mount immunostaining of DRGs for CD45, a pan-leukocyte marker, and podoplanin and Lyve-1, lymphatic endothelial markers used to define the meningeal surface. After PNI, the ipsilateral DRG meninges showed clustering of CD45^+^ cells, compared to sham and naïve controls (Fig. 1a-b; Fig. S1a-c). The CD45^+^ cell clusters were closely associated with the meninges and lymphatics surrounding the DRG, adhering to the DRG surface. The clusters were classified as medium and large, based on surface area statistics using the one-dimensional classification module in Imaris followed by automated surface segmentation. The DRG ipsilateral to PNI exhibited significantly greater numbers of CD45^+^ cell clusters across all size categories when compared to DRGs from sham-treated animals from both sexes (Fig. 1c-e). Representative 3D images depicting the medium and large clusters are shown in Fig. S2a. There were some CD45^+^ cell clusters in contralateral DRGs, though there were no gross differences between PNI or sham, and the frequency and size of these clusters were much lower compared to the ipsilateral DRG after PNI (Fig. S3a-c). The intensity of podoplanin and Lyve-1 was higher in PNI DRGs than in sham (Fig. 1f; Fig. S1d). The CD45^+^ cell clusters in the DRG meninges ipsilateral to PNI also contained MECA-79^+^ high endothelial venules (HEVs), which were absent after sham surgery (Fig. 1g-h). These specialized postcapillary venules allow lymphocyte transmigration into lymphoid organs, including TLSs^17, 18, 19, 21^.

**Figure 1.**
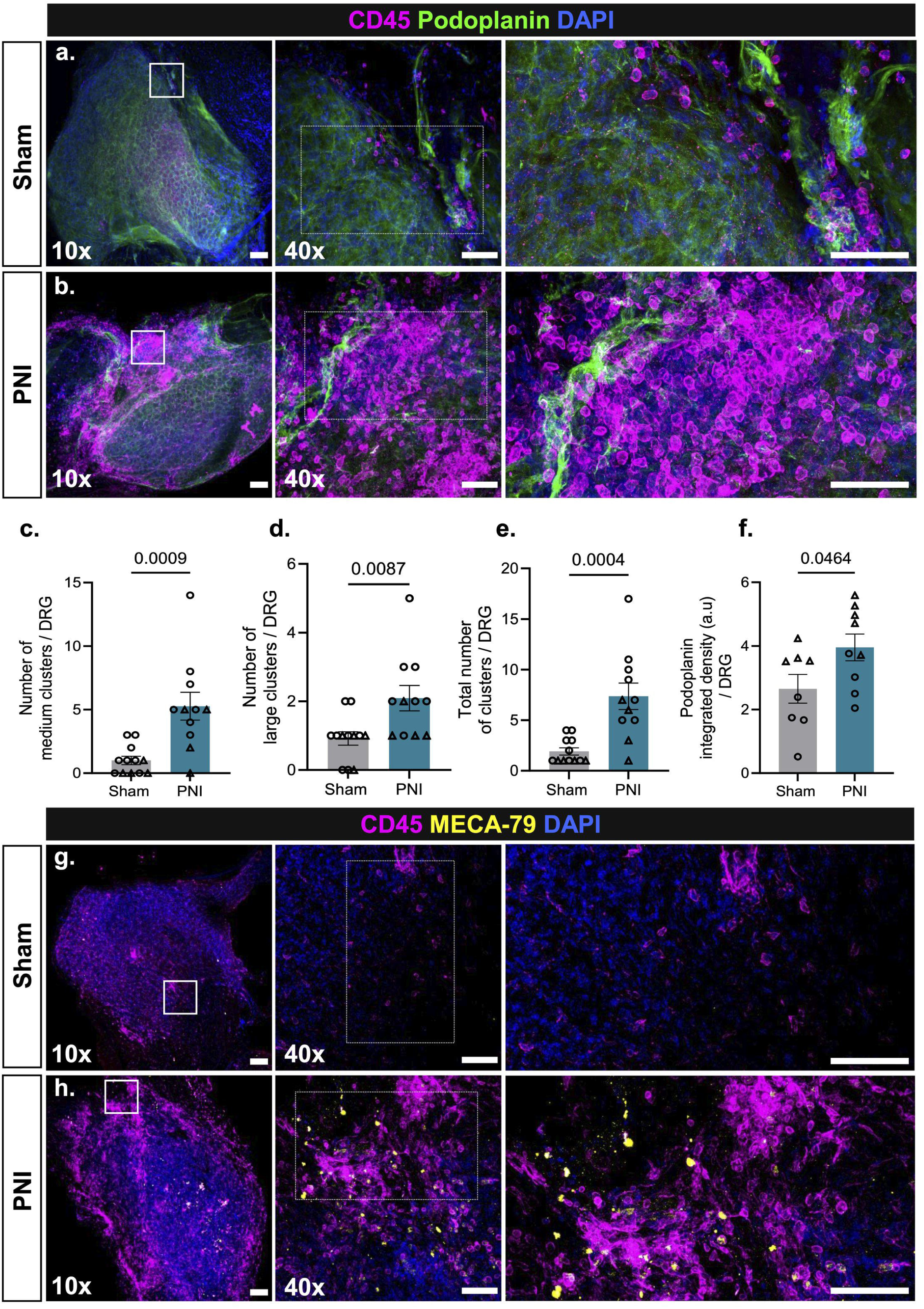
Meningeal CD45^+^ cell clusters develop in the ipsilateral DRG after PNI. (**a-b**) Representative 10x and 40x (inset) images of whole mount L4-L5 DRGs collected 14 days after PNI, and sham surgery (scale bar = 100 µm). The enlarged images were digitally cropped from the original 40x images (some are rotated, scale bar = 50 µm). DRGs were stained for a pan-leukocyte marker (CD45), lymphatic endothelial marker to define the meningeal surface (podoplanin), and a nuclear marker (DAPI) (**a**) ipsilateral to sham surgery (**b**) ipsilateral to PNI, (**c**) Numbers of medium, (**d**) large clusters, and total clusters (**e**) around DRGs from male and female mice were quantified. N = 11-12 per group. (**f**) Integrated density of podoplanin. N = 8-9 per group. Data analyzed by unpaired, two-tailed Student’s t tests. **(g-h)** Representative 10x, 40x (inset), and enlarged images (from 40x) displaying high endothelial venules (MECA-79) in close proximity to CD45^+^ cell clusters around the DRG after PNI.

We next sought to characterize the cellular composition of these clusters in ipsilateral DRG meninges after PNI or sham surgery. PNI induced clustering of innate immune cells, enwrapped by podoplanin^+^ lymphatic endothelium of the meninges, comprising Ly6G^+^ neutrophils (Fig. 2a-c) and CD11c^+^ dendritic cells (Fig. 2d-f), as compared to sham DRG. Some of these dendritic cells were CD45^+^Gr-1^+^CD21/35^+^ follicular dendritic cells (Fig. 2g-i). We also phenotyped the lymphoid cells within these meningeal clusters. After PNI, CD3^+^ T cells, and CD3^+^BCL6^+^ and CD3^+^PD-1^+^ Tfh cells were present within the leukocyte clusters, but absent in sham controls (Fig. 2j-m; Fig. S4a-d). B220^+^ B cells, enveloped by podoplanin^+^ lymphatic endothelium, formed clusters in the meninges after PNI (Fig. 2n-p). B220^+^ B cells had a scattered distribution as single cells in sham-operated mice (Fig. 2n). Moreover, some of these B220^+^ B cells expressed the proliferation marker proliferating cell nuclear antigen (PCNA), otherwise absent in the meninges of sham-operated mice (Fig. 2q-s). We also determined that B220^+^ B cells were in close association with Ly6G^+^ neutrophils (Fig S5a-b), CD11c^+^ dendritic cells (Fig. S5c-d), F4/80^+^ macrophages (Fig. S5e-g), CD3^+^ T cells (Fig. S5h-i). Assessing a cross-section of the decalcified spinal column as an alternative approach, we also identified CD45^+^ clusters comprising B220^+^ B cells at the meningeal interface between the ipsilateral DRG and spinal cord, as well as the sciatic nerve (Fig. S6a-d). Many of the B cells in the clusters were B220⁺BCL6⁺ and B220⁺CD95⁺ germinal center (GC) B cells, increasing in number after PNI compared to sham GC-B cells (Fig. 3a-c; Fig. S7a-g). Furthermore, B220⁺CD138⁺ plasma cells were identified in close association with the meninges in PNI DRGs, but absent in sham DRGs (Fig. 3d-g). Flow cytometric analysis of DRG and meninges confirmed increased B cells (CD45⁺B220⁺) (Fig. 4a-b), proliferating/cycling B cells (B220⁺CCND3⁺) (Fig. 4c-d), GC B cells (B220⁺BCL6⁺, B220⁺CD95⁺) (Fig. 4e-h), and plasma cells (B220⁺CD138⁺) (Fig. 4i-j) after PNI, compared to sham. CD45^+^ cells (Fig. S8a-b), CD3⁺ T cells (Fig. S8c-d) and CD3⁺BCL6⁺ Tfh cells (Fig. S8e-f) were similarly increased after PNI. These data demonstrate that PNI induces clustering of leukocytes in the DRG meninges, closely apposed by lymphatic vessels and HEVs which facilitate leukocyte trafficking. These meningeal B cell clusters contain the hallmark cellular constituents of germinal centers, including follicular dendritic cells, Tfh cells, and GC B cells, alongside proliferating B cells and plasma cells, consistent with bona fide TLSs.

**Figure 2.**
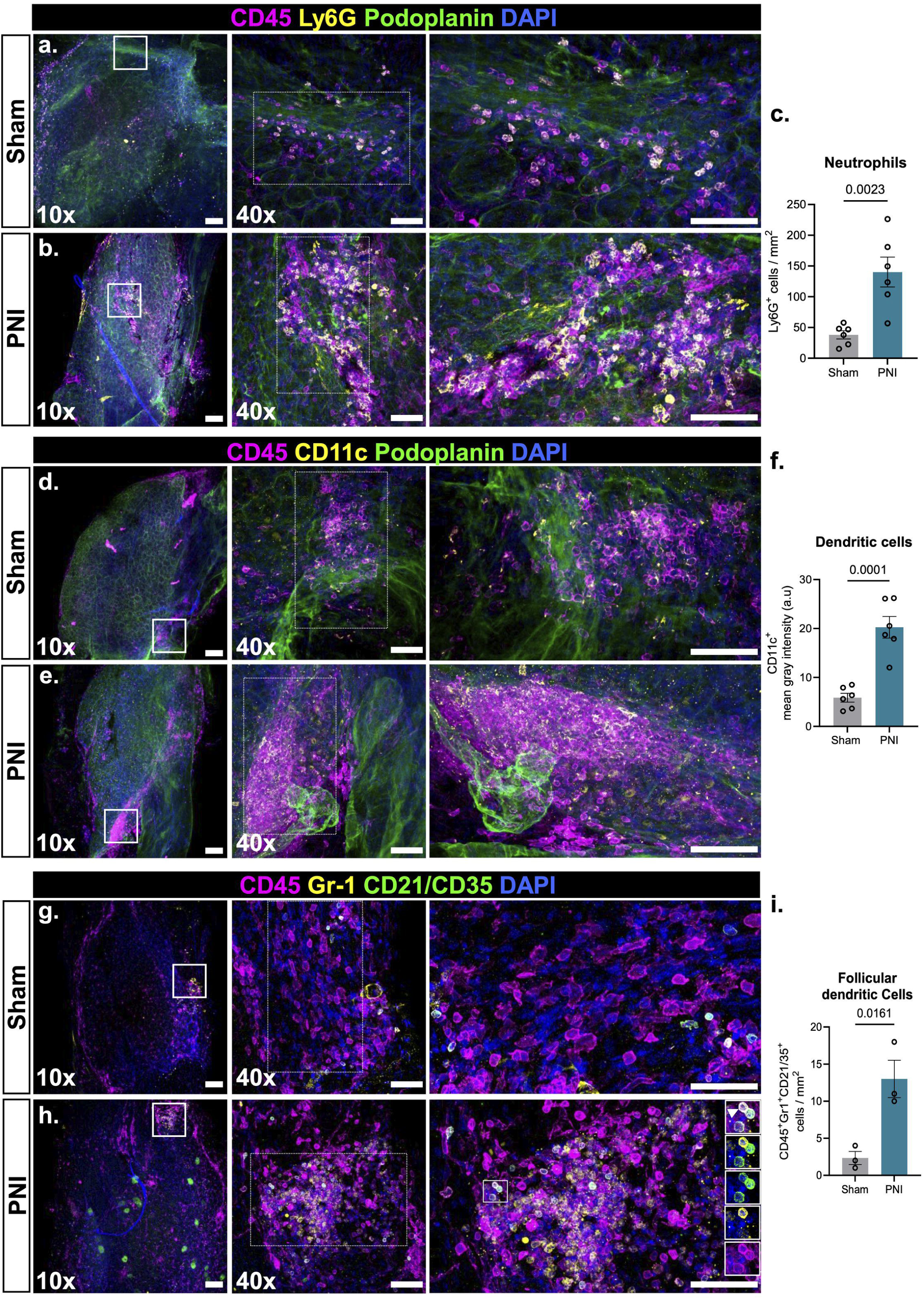
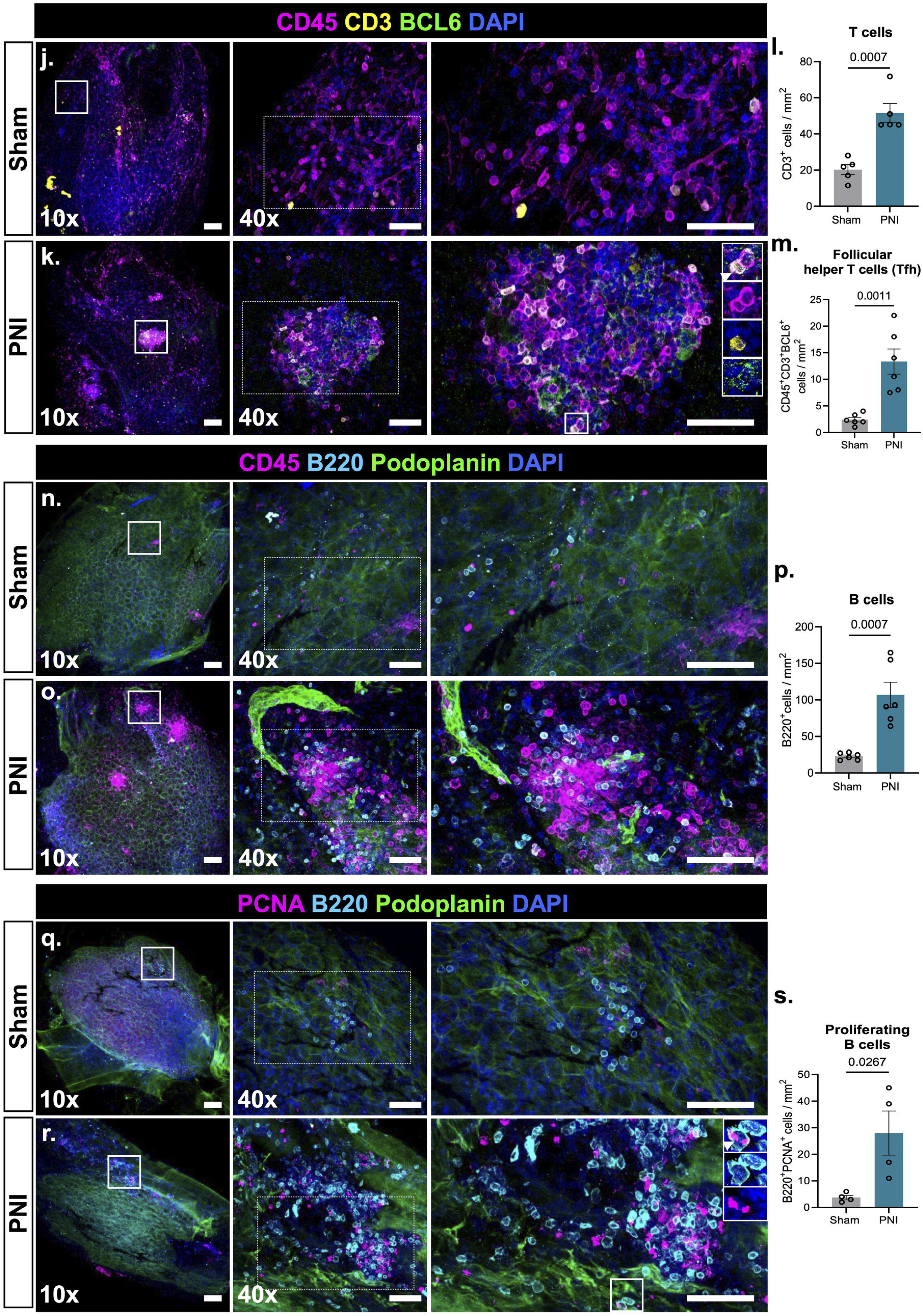
Clusters in DRG meninges are composed of B cells, T cells, neutrophils, Tfh cells and dendritic cells. Representative 10x and 40x (inset) images of whole mount ipsilateral L4-L5 DRGs collected 14 days after PNI or sham surgery (scale bar = 100 µm). The enlarged images were digitally cropped from the original 40x images (some are rotated, scale bar = 50 µm). (**a-b**) DRGs immunostained for neutrophils (Ly6G), a pan-leukocyte marker (CD45), lymphatic endothelial cells (podoplanin), and DAPI. (**c**) The numbers of Ly6G^+^ neutrophils were quantified and analyzed by unpaired, two-tailed Student’s t test. N = 6 per group. (**d-e**) DRGs immunostained for dendritic cells (CD11c), a pan-leukocyte marker (CD45), podoplanin^+^ lymphatic endothelium, and DAPI. (**f**) Fluorescence intensity for CD11c^+^ dendritic cells was quantified and analyzed by unpaired, two-tailed Student’s t test. N = 6 per group. (**g-h**) DRGs immunostained for pan-leukocyte marker (CD45) with Gr-1, CD21/CD35, and DAPI to identify follicular dendritic cells (FDC) and the number of FDCs were quantified and analyzed by unpaired, two-tailed Student’s t test. N = 3 per group (**i**). The highlighted insets represent the FDCs (white arrows). (**j-k**) DRGs immunostained for CD45^+^CD3^+^BCL6^+^ T cells (Tfh cells), and DAPI. (**l-m**) The numbers of CD3^+^ T and CD45^+^CD3^+^BCL6^+^ (Tfh) cells were quantified and analyzed by unpaired, two-tailed Student’s t test. N = 5-6 per group. (**n-o**) DRGs immunostained for a pan-leukocyte marker (CD45), B220^+^ B cells, podoplanin^+^ lymphatic endothelium, and DAPI. (**p**) The numbers of B220^+^ B cells were quantified and analyzed by unpaired, two-tailed Student’s t test. N = 6 per group. (**q-r**) DRGs immunostained for B220^+^ B cells, proliferation marker (PCNA), podoplanin^+^ lymphatic endothelium, and DAPI. (**s**) B220^+^PCNA^+^ cells were quantified and analyzed by unpaired, two-tailed Student’s t test. N = 4 per group.

**Figure 3:**
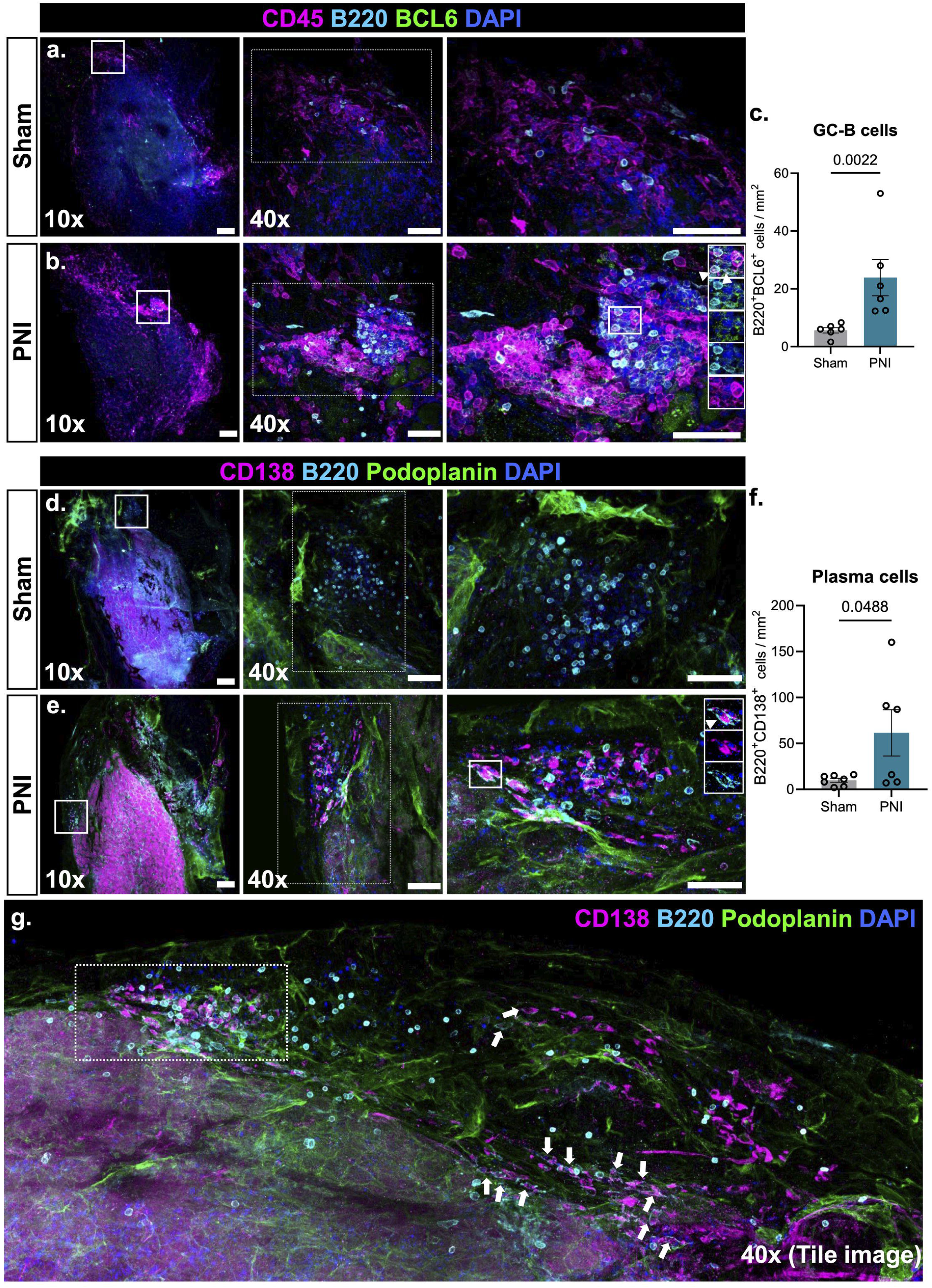
Germinal center B cells and plasma cell maturation are identified in DRG meninges after PNI. Representative 10x images with corresponding 40x inset images of wholemount ipsilateral L4–L5 DRGs collected 14 days after PNI or sham surgery (scale bar = 100 µm). Enlarged inset images were digitally cropped from the original 40x images (some rotated for presentation; scale bar = 50 µm). (**a–b**) DRGs were immunostained for the pan-leukocyte marker (CD45) with B cells (B220), the germinal center marker BCL6, and DAPI (nuclei). Highlighted insets indicate CD45⁺B220⁺BCL6⁺ cells (white arrows). (**c**) The numbers of B220⁺BCL6⁺ germinal center B cells were quantified and analyzed by unpaired, two-tailed Student’s t-test. N = 6 per group. (**d–e**) DRGs were immunostained for B cells (B220), plasma cells (CD138), lymphatic endothelial cells (podoplanin), and DAPI (nuclei). Highlighted insets indicate B220⁺CD138⁺ plasma cells (white arrows). (**f**) B220⁺CD138⁺ plasma cells were quantified and analyzed by unpaired, two-tailed Student’s t-test. N = 6 per group. (**g**) Representative 40x tile-scan image of wholemount DRGs from PNI mice stained for B220^+^, CD138^+^, podoplanin, and DAPI, showing the distribution of plasma cells across the DRG meninges.

**Figure 4:**
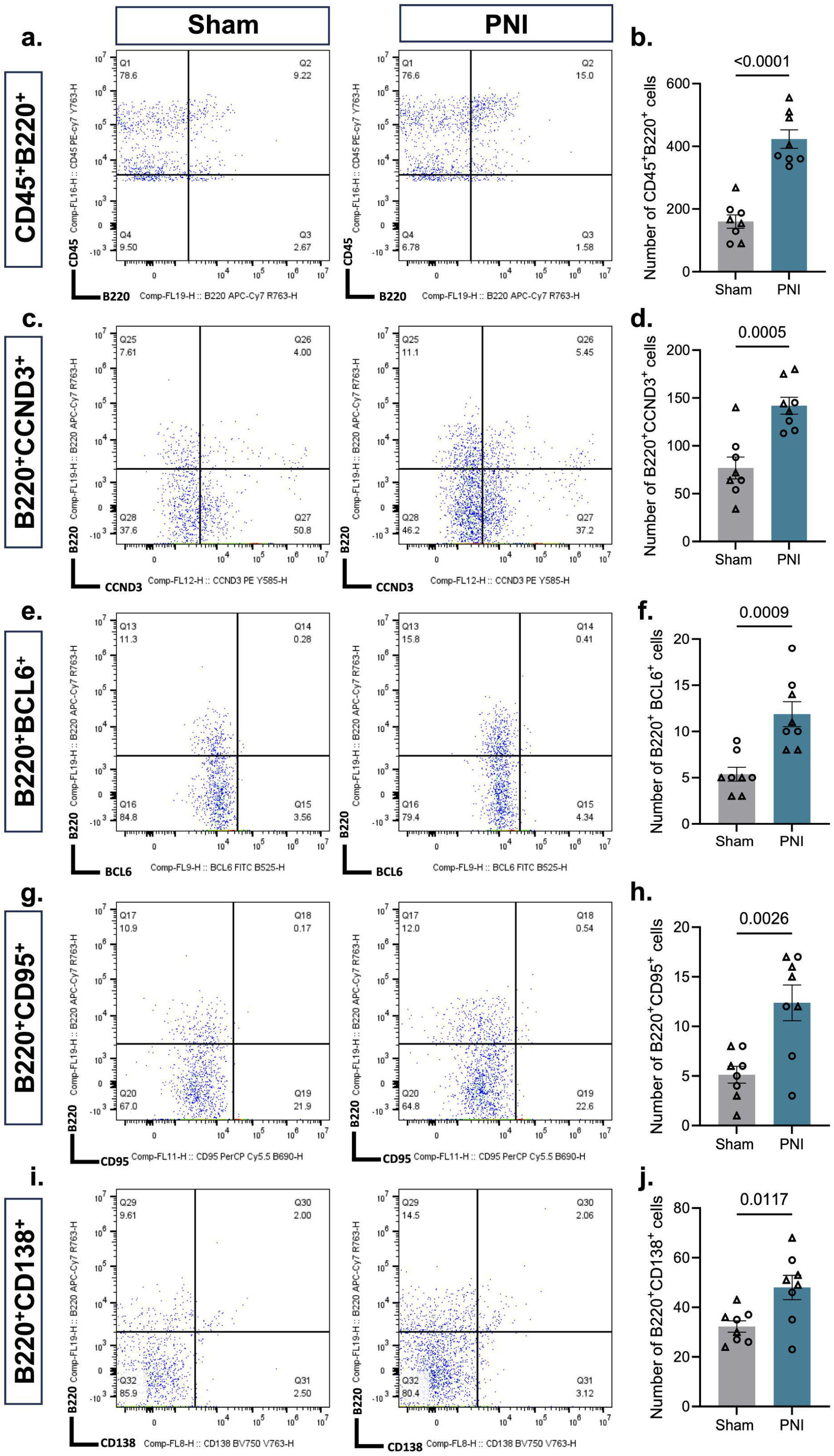
Flow cytometric validation of germinal center-associated B cell signatures in DRGs after PNI. DRGs with attached meninges were collected from PNI and sham mice 14 days after surgery and processed for flow cytometric analysis. Live CD45⁺ immune cells were gated and subsequently analyzed for B cell and germinal center-associated markers. (**a–b**) Representative scatter plots and quantification of the number of CD45⁺B220**⁺** B cells in sham and PNI groups. (**c–d**) Representative scatter plots and quantification of the number of B220⁺CCND3⁺ activated/proliferating B cells in sham and PNI groups. (**e-f**) Representative scatter plots and quantification of the number of B220⁺BCL6⁺ germinal center B cells in sham and PNI groups. (**g–h**) Representative scatter plots and quantification of the number of B220⁺CD95⁺ germinal center-associated B cells in sham and PNI groups. (**i-j**) Representative scatter plots and quantification of the number of B220⁺CD138⁺ plasma cells in sham and PNI groups. Each biological replicate consisted of pooled L3–L5 DRGs with attached meninges from three mice. n = 8 biological replicates per group (24 mice per group; 12 males and 12 females). Data was analyzed using unpaired, two-tailed Student’s t-test.

### Enriched transcriptional signature for meningeal germinal center B cells after PNI

To characterize molecular changes in dorsal root ganglia meningeal B cells after PNI (spared nerve injury, second model of PNI), we performed single-cell RNA-sequencing of CD45^+^ sorted cells from the DRG and dorsal root meninges. Eight main clusters of B cells were revealed through unsupervised clustering with t-distributed stochastic neighbor embedding (tSNE) (Fig. 5a-b). These clusters were assigned to known B cell subsets by comparing differentially expressed genes with established landmark genes (Fig. S9a). Among the meningeal B cells obtained from naïve mice (2268 cells), we identified immature B cells, naïve and mature follicular B cells, antigen-presenting cell (APC) like B cells, activated B cells and mature cells (Fig. 5a-c). From PNI mice (776 cells), we found mainly pre-GC and GC B cells (Fig. 5b-c). The dominant transcriptional signature for germinal center B cells is consistent with TLS development after PNI.

**Figure 5.**
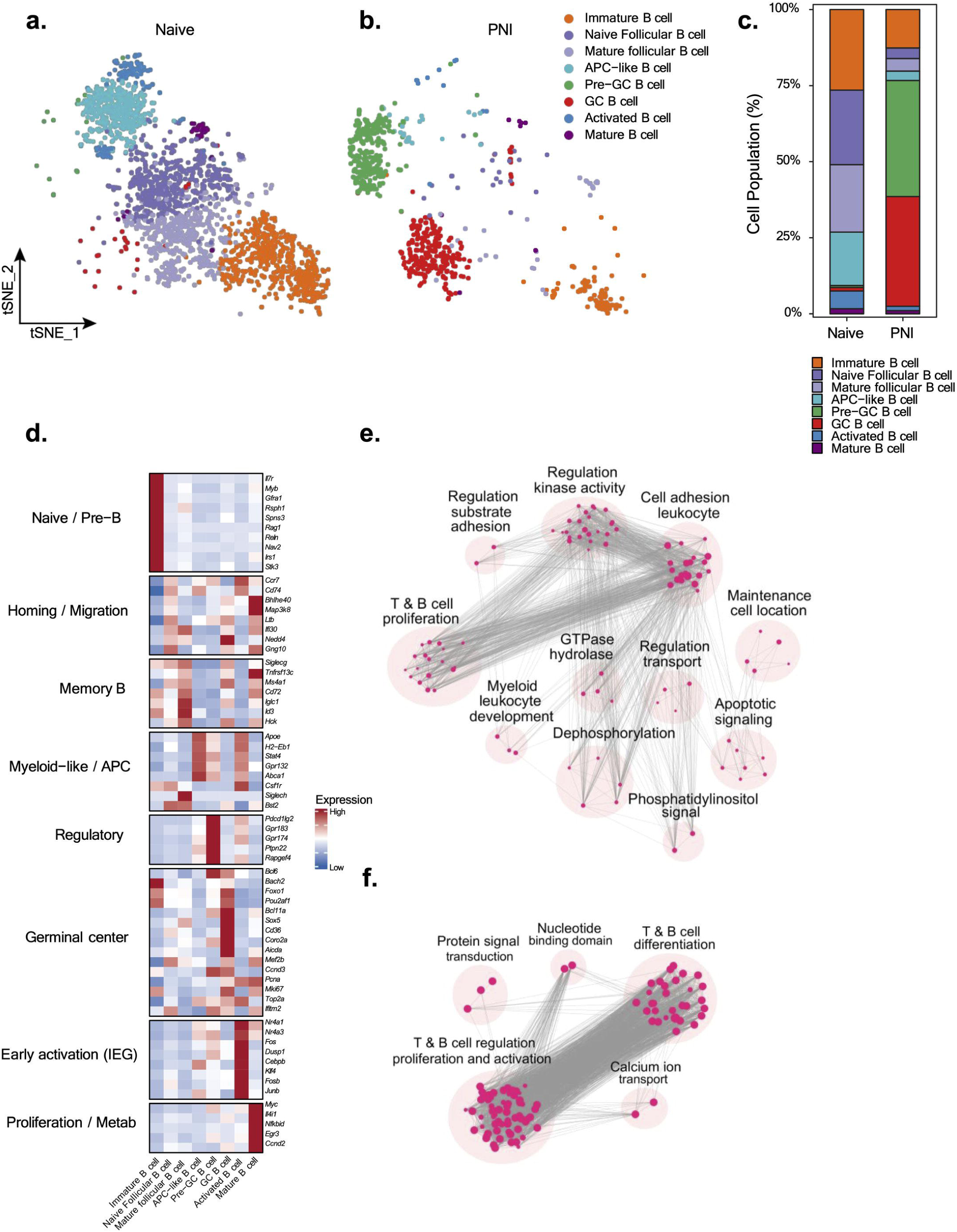
Enriched transcriptional signature for meningeal germinal center B cells after PNI. (**a-b**) Clustering of B cells identified from single-cell RNA sequencing of CD45^+^ cells in the dorsal root ganglia meninges from naïve mice and PNI mice (spared nerve injury, second model of PNI). Clustering of B cell subsets in naïve (360 cells) and PNI mice (776 cells). (**c**) Percentage of B cell subsets relative to the whole B cell population. (**d**)Heatmaps of genes differentially expressed between germinal center B cells and other subsets. (**e-f**) Pathway network analysis of upregulated pathways in GC and pre-GC clusters.

We assessed the genes which were differentially regulated between GC B cells and other subsets (Fig. 5d). GC B cells were distinguished from all other subsets by differential expression of genes involved in GC formation, maintenance, somatic hypermutation and affinity maturation (*Bcl6, Foxo1, Pou2af1, Aicda, Mef2b, Bcl11a).* Additional enrichment of *Sox5, Cd36, Coro2a, Top2a, Ifitm2* suggests contributions to GC B cell differentiation, metabolic adaptation, proliferation, and immune activation. These clusters also exhibited elevated expression of cell cycle associated genes *Ccnd3, Pcna,* and *Mki6* indicating a highly proliferative GC-B cell population. The pre-GC population was enriched with *Bcl6, Ccnd3, Bcl11a, Coro2a, Ifitm2* indicating acquisition of an early GC transcriptional program. This population also expressed regulatory and migration-associated genes, including *Pdcd1lg2, Gpr183, Gpr174, Ptpn22,* and *Rapgef4*, suggesting a transitional state preceding full GC differentiation.

We further analyzed the top enriched pathways associated with the GC and pre-GC B cell populations, represented as a pathway network map (Fig. 5e-f). The major pathways were related to B cell differentiation and activation (*Bcl11a, St3gal1, Hdac9, Card11, Ptprj, Ptprc, Rasgrp1, Btla, Card11, Cd180*) indicating active regulation of B cell maturation and signaling. In addition, pathways associated with positive regulation of T cell activation and differentiation were identified (*Rasgrp1, Dusp10, Card11, Hsph1, Cd274, Bcl11a, Egr1, Cdk6, Ptprc, Vav1*), suggesting coordinated B–T cell interactions within the GC microenvironment. Enrichment of genes involved in regulation of leukocyte cell-cell adhesion (*Rasgrp1, Btla, Dusp10, Card11, Hsph1, Ptprc, Vav1, Cd274, Pag1*) were also identified, indicating enhanced immune cell communication and cellular organization during GC formation and maturation.

### DRG meningeal TLSs are conserved in pigs with PNI and present in humans with chronic pain

Having established that PNI drives the formation of TLSs in the mouse DRG meninges, we next asked whether this response is conserved across mammalian species and relevant to human pain pathology. We first examined DRGs from tail-docked piglets and corresponding control animals. Immunostaining of cryosectioned DRGs revealed increased IgG deposition and CD79a^+^ B cell intensity in tail docked DRGs compared with controls (Fig. 6a-d). Wholemount immunostaining further identified BA4D5^+^ macrophages and CD79a^+^ B cell aggregation forming TLSs within PNI DRGs (Fig. 6e-f).

**Figure 6:**
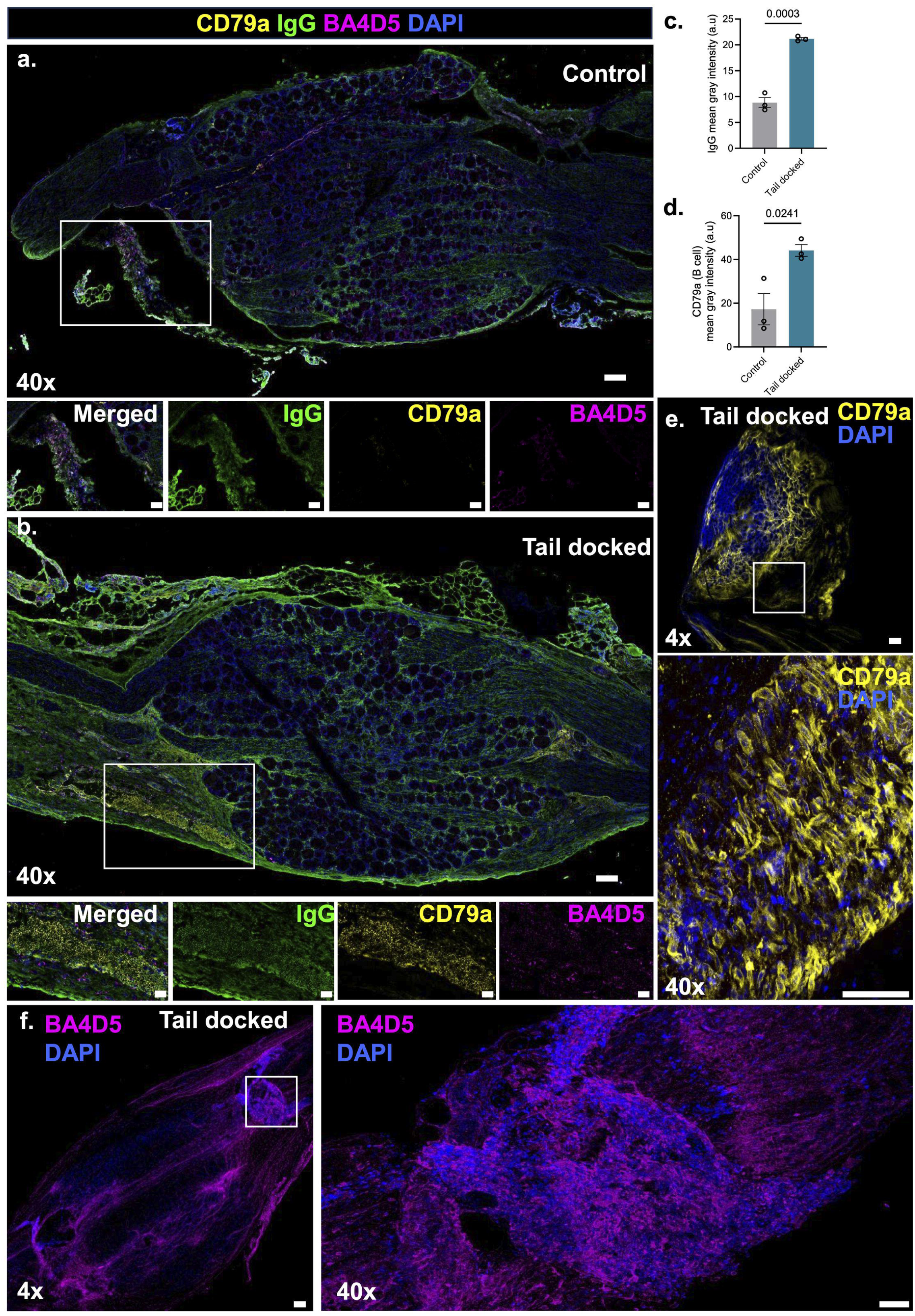
Immune clusters are identified in DRGs of tail docked piglets. (**a–b**) Representative 40x tile-scan images of cryosectioned ipsilateral L5 DRGs stained for CD79a (B cells), IgG, and BA4D5 (macrophages) from piglets collected 14 days after tail docking or control surgery (scale bar = 100 µm). (**c–d**) Mean gray intensity of IgG and CD79a staining was quantified and analyzed using an unpaired, two-tailed Student’s t-test. N = 3 animals per group. (**e–f**) Representative 4× wholemount images (scale bar = 100 µm) with corresponding 40x inset images (scale bar = 50 µm) of L4–L5 DRGs from tail-docked piglets stained with CD79a and DAPI (**e**), or BA4D5 and DAPI (**f**), demonstrating TLS-like immune clusters within intact tissue.

We next sought to determine whether TLSs are present in the human DRG (hDRG) in the context of pain pathology. We reanalyzed our published bulk RNA-sequencing dataset from surgical hDRG samples recovered from individuals with or without neuropathic pain^9^, examining two gene sets: one derived from GC B cell, Tfh cell, and follicular dendritic cell marker genes identified in this study (*TLS_curated gene set*), and a second from a published TLS signature in human cancers (*TLS_Meylan2022*)^22^. Although variability across samples suggested that TLS presence may be limited to a subset of neuropathic pain patients, analysis of the top quartile revealed substantial enrichment of TLS-associated genes in most pain-associated samples using both gene sets (Fig. 7a-b). Clonotype analysis of this cohort identified outliers with high BCR and TCR clonotype counts selectively in the pain group (Fig. 7c-d). To spatially resolve TLS architecture at the cellular level, we examined the hDRG of a representative organ donor with severe lumbar disc degeneration and a high DRG B cell transcriptional signature using imaging-based spatial transcriptomics (Fig. 7e). A TLS was identified within the DRG parenchyma, located within 50 µm of sensory neuron cell bodies (Fig. 7f). RNA transcript distribution and unsupervised clustering revealed an unusually large vessel cross-section demarcated by a ring of endothelial cells and surrounded by distinct B cell and T cell zones within an aggregation of mononuclear phagocytes (Fig. 7g-h). Smaller lymphoid structures were observed near microvasculature throughout the tissue section, with B cells identifiable by their unusually high interior RNA content and T cells noted to breach the satellite glial cell envelope (Fig. 7i-j). BCR and TCR clonotype analysis from matched bulk RNA-seq detected 221 unique clonotypes, the majority derived from BCRs, confirming the functional maturity of the TLS (Fig. 7k). Together, these findings demonstrate that injury-associated TLS formation in the DRG is conserved across mammalian species and that TLSs develop within the hDRG parenchyma in proximity to sensory neurons, where they are associated with pain pathology in a subset of patients.

**Figure 7.**
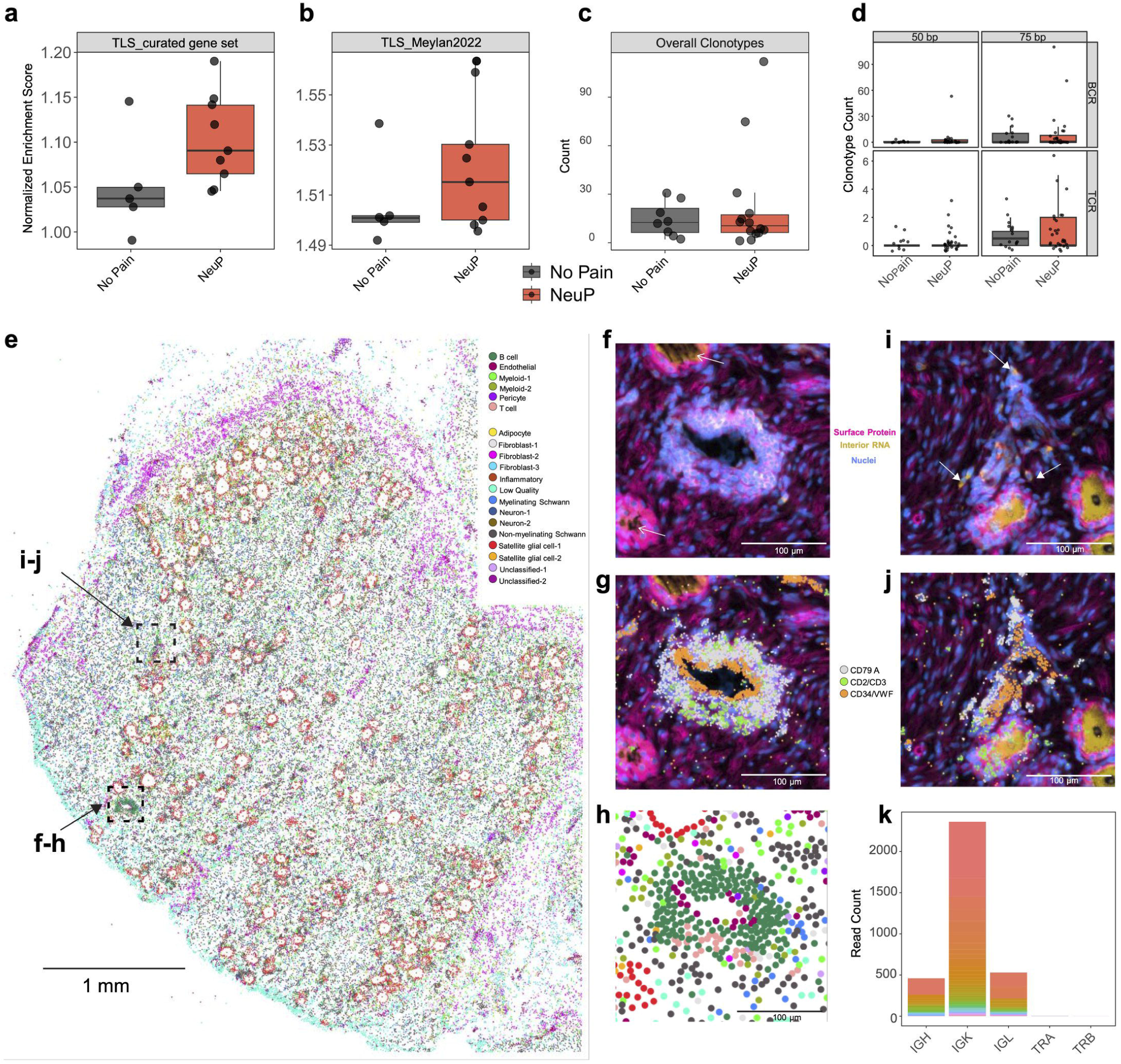
Spatial transcriptomics and bulk RNA sequencing of human DRGs identify TLS signatures and germinal center-associated markers. Tertiary lymphoid structures can be found in human dorsal root ganglion (DRG) parenchyma and are associated with pain. (**a-b**) Assessment of previously published RNA sequencing data for TLS-associated genes in human DRGs sourced from patients with and without neuropathic pain. In two unique TLS-associated gene sets (*TLS_curated gene set* and *TLS_Meylan2022*), the top quartile of genes were enriched in DRGs from donors with neuropathic pain (NeuP) compared to those with no pain. (**c**) DRGs from donors without neuropathic pain had low (<30) unique BCR clonotypes detectable in bulk RNA sequencing (single-end 75 base pair samples). Two outliers were found in the neuropathic pain group with high BCR clonotype diversity. (**d**) Clonotype analysis of previously published dorsal root ganglia bulk RNA sequencing derived from human donors with and without neuropathic pain. Data generated from^49^. Frequency of unique clonotypes mapped to B cell receptors (BCR) or T cell receptors (TCR). Samples sequenced with a single-end read length of 50 or 75 base pairs were plotted separately. Outliers with high numbers of unique clonotypes were found in samples with both read lengths, but the 75 bp read length generated a more sensitive and suitable MiXCR analysis. (**e**) A human DRG tissue section from a donor with severe lumbar spine degeneration shows a TLS and other sites of B cell aggregation. Each dot is a cell centroid based on clustering against 472 genes probed in 10x Xenium imaging-based spatial transcriptomics. (**f**) TLS histology shows a dense area of cells surrounding a blood vessel cross-section within 50 µm of sensory neuron cell bodies (open arrowheads). (**g**) RNA transcripts marking B cells (*CD79A*), T cells (*CD2*, *CD3D*, *CD3E*, *CD3G*), and endothelial cells (*CD34*, *VWF*) demonstrate a central ring of endothelial cells, multiple surrounding B cells, and a distinct distal T cell zone interacting with the perivascular B cells. (**h**) Magnified TLS transcriptomics from (e) confirm B cell (dark green) and T cell (light pink) identity and further reveal two distinct mononuclear phagocyte subpopulations (lime, olive) interacting with the B cell and T cell zones of the structure. (**i**) B cells are further distributed throughout this DRG and are distinguishable by high intensity 18S interior RNA staining (closed arrowheads). (**j**) These B cells are adjacent to blood vessels nearby sensory neurons and T cells that breach the satellite glial cell envelope. (**k**) Bulk total RNA sequencing taken from additional tissue sections from this DRG detected thousands of reads from 221 unique BCR and TCR clonotypes (*IGH*, *IGK*, and *IGL* family genes corresponding to BCR heavy and light chains, and *TRA* and *TRB* family genes corresponding to TCR alpha and beta chains).

### B cells in the lumbar region facilitate mechanical allodynia after PNI

The dominant presence of TLSs in the DRG meninges after PNI led us to consider whether B cells within these structures contribute to pain. Our previous work using systemic depletion or deficiency approaches demonstrated that B cells are required for the development of mechanical hypersensitivity (allodynia) after PNI in mice^9^. Here, we tested whether allodynia is influenced by B cells in the lumbar region, including TLSs in the DRG meninges. Eliminating B cells with a single intrathecal injection of CD20 monoclonal antibody (mAb) at the time of PNI prevented development of mechanical allodynia for at least four weeks (Fig. 8a). In contrast, injured mice treated with isotype control mAb developed mechanical allodynia by day 7 post-PNI, which was sustained through four weeks after injury (Fig. 8a). CD20 mAb treatment of sham-operated mice did not alter paw withdrawal thresholds (Fig. 8a).

**Figure 8.**
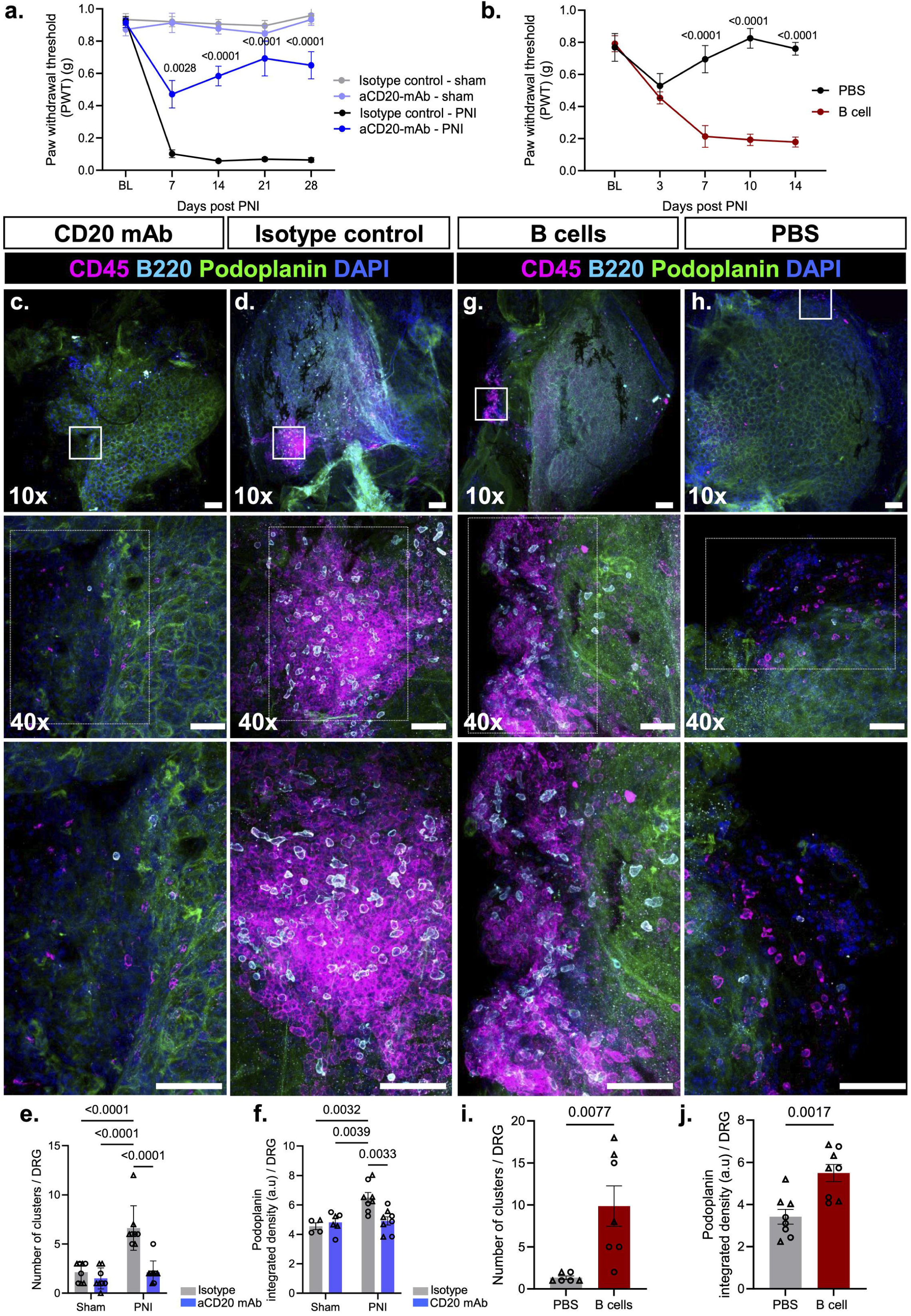
B cells in the lumbar region are required for mechanical allodynia and TLS formation after PNI. (**a**) Von Frey thresholds for mechanical allodynia were assessed for ipsilateral hindpaws in both male and female mice that received a single intrathecal injection of CD20 mAb or IgG isotype control antibody (10 μg) at the time of PNI/sham surgery. Data analyzed by repeated measures two-way ANOVA and Sîdak’s post hoc tests. N = 7-8 males per group and 3-5 females per group. Baseline (BL) measures before surgery/injections. (**b**) Von Frey thresholds were assessed for ipsilateral hindpaws of muMT male and female mice receiving intrathecal injections of B cells (2 x 10^5^ in 5 μl) or PBS control immediately before PNI. Data analyzed by repeated measures two-way ANOVA and Sîdak’s post hoc tests. N = 3-5 males per group and 3 females per group. (**c-d**) Representative 10x and 40x (inset) images of whole mount ipsilateral L4-L5 DRGs collected 28 days after PNI/sham surgery or intrathecal CD20 mAb/IgG isotype control antibody treatment (scale bar = 100 µm). The enlarged images were digitally cropped from the original 40x images (scale bar = 50 µm). DRGs obtained from mice after PNI and treatment with CD20 mAb or isotype control were immunostained for B cells (B220), pan-leukocytes (CD45), lymphatic endothelial cells (podoplanin), and DAPI (nuclei). (**e**) Total numbers of medium + large clusters around male and female DRGs were quantified. N = 8-9 per group. Data analyzed by two-way ANOVA and Tukey’s multiple-comparisons test. (**f**) Integrated density of podoplanin intensity was measured data analyzed by two-way ANOVA and Tukey’s multiple-comparisons test. N = 4-8 per group (**g-h**) Representative 10x and 40x (inset) images of whole mount ipsilateral L4-L5 DRGs collected 14 days after PNI/sham surgery or intrathecal B cells/PBS control treatment (scale bar = 100 µm). The enlarged images were digitally cropped from the original 40x images (some are rotated, scale bar = 50 µm). DRGs obtained after PNI and B cell or PBS injection were immunostained for B cells (B220), pan-leukocytes (CD45), lymphatic endothelial cells (podoplanin), and DAPI (nuclei). **(i**) Total numbers of medium + large clusters in male and female DRGs were quantified. N = 6-7 per group. Data analyzed by unpaired, two-tailed Student’s t tests. **(j)** Integrated density of podoplanin intensity was measured and data analyzed by unpaired, two-tailed Student’s t tests. N = 8 per group.

To confirm that intrathecal CD20 mAb did not influence systemic antibody secretion, we assessed the secondary IgG response 14 days after tetanus toxoid booster injections (which occurred 7 days after CD20 mAb treatment). Equivalent levels of serum anti-tetanus toxoid IgG were observed following intrathecal CD20 mAb or IgG isotype control (Fig. S10a). These results confirm that this intrathecal CD20 mAb treatment left systemic adaptive immune responses intact.

As a complementary approach, we tested whether B cells in the lumbar region were sufficient for induction of allodynia after PNI. Purified splenic B cells from wild-type mice (or phosphate buffered saline (PBS) control) were intrathecally injected into B cell-deficient mice (muMT mice) at the time of PNI. Whereas PBS-treated muMT mice did not develop mechanical hypersensitivity, allodynia was reinstated in these mice by transfer of B cells into the lumbar region (Fig. 8b). These results collectively demonstrate that lumbar B cells are essential for the development of mechanical allodynia in mice after PNI.

### B cells in the lumbar region are required for TLS organization in the meninges of the DRG

Having established that lumbar B cells drive allodynia, we next turned to the structural mechanism underlying this effect. We assessed the influence of single dose intrathecal CD20 mAb on B cell numbers and TLS organization in the DRG. Strikingly, CD20 mAb treatment not only eliminated meningeal B220^+^ B cells present in the clusters after PNI, but virtually all CD45^+^ cell clusters, compared with isotype control (Fig. 8c-e). Consistently, localized B cell depletion also reduced IgG accumulation in cryosectioned DRGs, compared with isotype control (Fig. S11a-b). Podoplanin intensity was reduced by CD20 mAb treatment, compared to control (Fig. 8f). No treatment-dependent differences in cluster formation were observed in sham DRGs (Fig. S11c-d). Flow cytometric analysis confirmed depletion of CD45⁺CD19⁺ and CD45^+^B220^+^ B cells in DRG, with no change in total CD45^+^ cell numbers (Fig. S12a-b).

Consistent with these findings, CD45^+^ cell clusters were absent from DRG meninges of muMT mice of both sexes after PNI (Fig. 8g-h). Intrathecal transfer of B cells to muMT mice at the time of PNI was sufficient for organization of CD45^+^ cell clusters, dominated by B220^+^ cells (Fig. 8i), and increased podoplanin intensity, compared with PBS controls (Fig. 8j). These data reveal that B cells in the lumbar region are necessary and sufficient for TLS organization in the DRG meninges.

### Germinal center B cells are required for TLS organization and allodynia after PNI

The absence of TLSs after B cell depletion raised the question of which subset(s) of B cells would regulate their organization. Given that our data identify enrichment of GC B cells, we tested the role of this subset directly. The EZH2 histone methyltransferase is preferentially expressed in germinal center B cells, where is facilitates rapid proliferation by repressing cell cycle checkpoint genes^23, 24, 25, 26, 27^. To restrict *Ezh2* deletion to B cells undergoing the GC response, we generated *Aicda*^creERT2/+^::*Ezh2*^fl/fl^ mice and evaluated TLS formation and pain behavior after PNI. Clusters of CD45^+^ cells and B220^+^ B cells were absent from the meninges of *Aicda*^creERT2/+^::*Ezh2*^fl/fl^ mice of both sexes, compared to littermate controls (Fig. 9a-c). There were no differences in podoplanin intensity between *Aicda*^creERT2/+^::*Ezh2*^fl/fl^ mice and littermate controls (Fig. 9d). No TLSs were identified in the sham DRGs (Fig. S13a-b). Moreover, *Aicda*^creERT2/+^::*Ezh2*^fl/fl^ mice did not develop mechanical allodynia for at least four weeks after injury, in contrast to littermate controls (Fig. 9e), while paw withdrawal thresholds of sham-operated mice were unaltered (Fig. 9e). These data demonstrate that germinal center B cells are required for both TLS organization and the development of mechanical allodynia after PNI.

**Figure 9.**
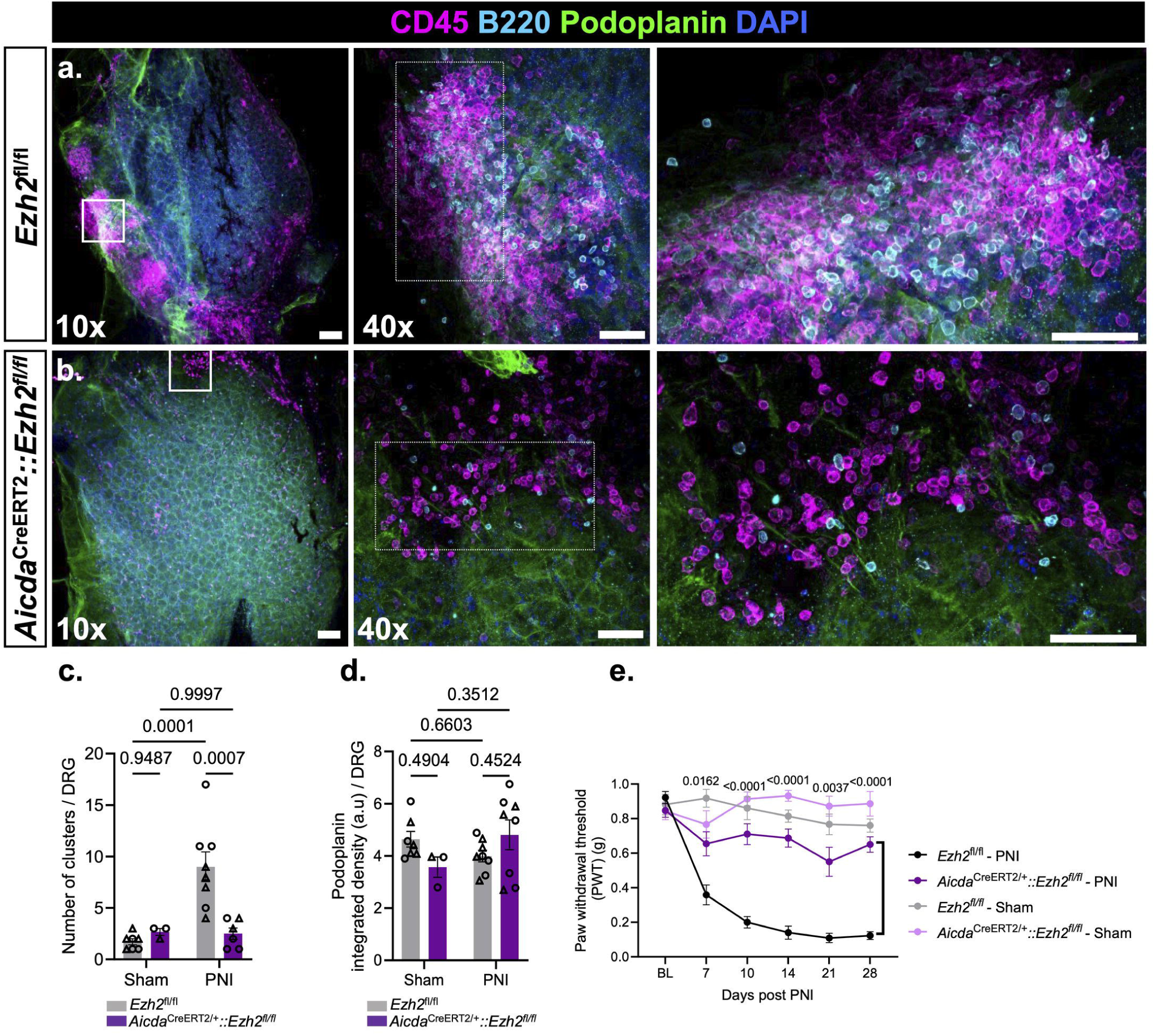
PNI does not induce TLS formation or mechanical allodynia in *Aicda*^creERT2/+^::*Ezh2*^fl/fl^ mice. (**a-b**) Representative 10x and 40x (inset) images of whole mount ipsilateral L4-L5 DRGs collected 28 days after PNI surgery in *Aicda*^creERT2/+^::*Ezh2*^fl/fl^ male mice or littermate control mice (scale bar = 100 µm). The enlarged images were digitally cropped from the original 40x images (some rotated for presentation; scale bar = 50 µm). DRGs were obtained from *Aicda*^creERT2/+^::*Ezh2*^fl/fl^ or littermate mice after PNI and immunostained for B cells (B220), pan-leukocytes (CD45), lymphatic endothelial cells (podoplanin), and DAPI (nuclei). (**c**) Numbers of medium + large clusters around male and female DRGs in each group. Data analyzed by two-way ANOVA and Tukey’s multiple-comparisons test. N = 3-8 per group. (**d**) Integrated density of podoplanin intensity was measured data analyzed by two-way ANOVA and Tukey’s multiple-comparisons test. N = 3-8 per group (**e**) Von Frey thresholds for mechanical allodynia were assessed for ipsilateral hindpaws of male and female *Aicda*^creERT2/+^::*Ezh2*^fl/fl^ mice or littermate control mice that received PNI or sham surgery. Data analyzed by repeated measures two-way ANOVA and Sîdak’s post hoc tests. N = 3-5 males and 2-5 females per group.

## DISCUSSION

We have discovered that PNI induces clustering of leukocytes apposed to lymphatic vessels and HEVs in the ipsilateral DRG meninges, consistent with TLSs. These structures are germinal center-like, composed of follicular dendritic cells and Tfh cells, as well as proliferating B cells, GC B cells, and plasma cells. Similar TLS-like immune clusters were identified in tail-docked piglets, and in human DRGs from donors with neuropathic pain, where spatial transcriptomics identified mature TLSs within the DRG parenchyma and BCR/TCR clonotype analysis confirmed their functional maturity. Transcriptional analyses identified enrichment of germinal center B cells in the dorsal root ganglia meninges after PNI. Germinal center B cells are pivotal for TLS organization: these structures were absent after PNI in mice with impaired B cell clonal expansion and somatic hypermutation due to *Ezh2* deletion^25, 26, 27^. Local depletion of B cells in the lumbar region with intrathecal CD20 mAb also prevented TLS organization. Conversely, intrathecal transfer of B cells to muMT mice was sufficient to restore TLS organization after PNI. These structures appear to be pronociceptive, as allodynia failed to develop when TLS organization was disrupted by intrathecal CD20 mAb, *Ezh2* deletion, or muMT genotype, and was reinstated in muMT mice by intrathecal B cell transfer at the time of PNI. Thus, B cell signaling from the lumbar region is crucial for TLS organization and mechanical allodynia induced by PNI.

The presence of TLSs is associated with a less favorable prognosis in autoimmune conditions like multiple sclerosis/experimental autoimmune encephalomyelitis, systemic lupus erythematosus, and arthritis^17, 18, 19^. One reason proposed is that these TLS are probably sites of autoantibody production. In our earlier work, we showed that autoreactive IgG production causes pain after PNI in mice^9, 10, 11^. Other groups have further implicated autoreactive antibodies in chronic musculoskeletal and neuropathic pain^28, 29, 30, 31, 32, 33, 34, 35, 36^. One could predict that the TLSs associated with the DRG meninges are local sites where novel autoantigens, released after nerve injury, are presented by dendritic cells/macrophages in the TLS to Tfh cells which activate B cells, leading to their differentiation into antibody-secreting cells. Unlike secondary lymphoid organs, TLSs require persistent antigenic and inflammatory stimulation to be maintained and typically regress once this drive is withdrawn^21^. Because the TLSs we describe form at a site — the DRG meninges — distal to the peripheral nerve injury itself, it is unclear how antigenic supply to them is sustained over time. Extracellular vesicles containing de novo autoantigenic epitopes derived from injured tissue could be one source^37^. These vesicles formed at the site of injury, can be transported by dendritic cells which subsequently home to the lymphoid organ (in this case the TLS), where their antigenic content can be processed and initiate the auto-immune response^37^. Antigen-loaded dendritic cells or macrophages that acquire autoantigen at the injury site could similarly traffic directly to the DRG meninges and re-present this cargo locally, sustaining the germinal center reaction^38^. Other routes are also plausible: damage-associated antigen taken up at the injured nerve terminal could undergo retrograde axonal transport back to the DRG soma and meninges, exploiting the unique anatomy of pseudounipolar sensory neurons^39, 40^; or antigen may not need to travel at all, if axotomized DRG neurons generate neoantigens locally as part of their own injury response^41^. Distinguishing among these routes will require dedicated antigen-tracing studies. It is of further interest that pain is alleviated by enhanced efferocytosis at the PNI site^42^, which might be partly due to less de novo autoantigen release at the site of injury.

We have observed follicular dendritic cells, Tfh cells, and HEVs, together with B cells expressing the germinal center markers BCL6 and CD95, and CD138^+^ plasma cells, indicating the formation of functional germinal centers, supporting our conclusion that TLSs are organized in the DRG meninges after PNI. Transcriptional analysis also revealed enrichment of germinal center B cells, including canonical genes governing germinal center formation and somatic hypermutation (e.g., *Bcl6*, *Aicda*, *Mef2b*) in the dorsal root ganglia meninges after PNI. With follicular dendritic cells, Tfh cells, germinal center (BCL6^+^/CD95^+^) B cells, and plasma cells all identified by immunostaining, flow cytometry, and single-cell transcriptomics, these clusters display the hallmark cellular constituents of a germinal center reaction, even though classic follicular segregation into dark and light zones, as seen in secondary lymphoid organs, has not yet been demonstrated. Cross-species relevance was further supported by tail-docked piglets, which showed increased IgG deposition co-localized with B cell aggregates in the DRG, and by BCR/TCR clonotype analysis of a human DRG TLS, which identified a restricted, predominantly B cell receptor-derived clonal repertoire indicative of a functionally mature, antigen-driven germinal center reaction. Our results demonstrate the necessity of the germinal center B cell subset (in which Ezh2 is deleted specifically from germinal center-experienced B cells) to support IgG as the critical effector for pain after PNI. We previously uncovered IgG deposition in the lumbar spinal cord and DRG of nerve-injured mice, and enrichment of a B cell transcriptional signature from DRG of human donors with neuropathic pain^9^. Moreover, we recently showed that pain after PNI was attenuated by locally depleting IgG around the lumbar region^10^. Autoreactive IgG forms complexes with their cognate neoantigens exposed and released after nerve injury^11^. These immune complexes can locally activate Fc gamma receptors to induce hyperexcitability of DRG neurons, leading to pain^9, 10, 11^. Here, we reveal DRG meningeal TLSs, which are dominated by B cells, as a locus for pronociceptive B cell signaling through IgG.

In addition to alleviating pain, depleting or interfering with lumbar B cell function disrupted TLS organization as well. The exact role of B cells in TLS formation or maintenance has not yet been established, but activated B cells are known to form follicle-like B structures with the support of Tfh and follicular dendritic cells^17, 18, 19^. B cell recruitment into these follicle-like aggregates is classically directed by the chemokine CXCL13, produced by stromal and follicular dendritic cells and sensed by CXCR5 on B cells^21^. Once established, cytokine production by the autoantigen-activated B cells could produce niche-promoting mediators like lymphotoxins (e.g., lymphotoxin-a/b) which could engage cognate receptors on stromal cells to promote TLS germinal centers^21, 43^. TLS assembly may also depend on relief of B cell-intrinsic inhibitory signaling, as autocrine lysophosphatidylserine normally restrains B cell aggregation, and its loss enhances germinal center and TLS-like formation^44^. Depletion of B cells, even with a single intrathecal treatment with CD20 mAb could therefore prevent TLS formation.

Leukocytes populating the meninges derive from the skull and vertebral bone marrow, migrating through osseous channels^45, 46, 47, 48^. In particular, the skull is a source of meningeal B cells in young mice, allowing for education by CNS antigens^45^. We have previously identified a resident population of B cells in the mouse and human DRG under naïve conditions^9^. It is possible that B cells (and other cells) comprising TLSs proliferate from a resident pool or are recruited from the vertebral bone marrow in response to PNI. Consistent with this route, peripheral nerve injury has recently been shown to drive emergency myelopoiesis in the vertebral bone marrow, with the resulting myeloid cells migrating through ossified vertebral channels into the DRG meninges under the instruction of meningeal GM-CSF from group 2 innate lymphoid cells ^12^; whether B cells reach the meninges via a similar route remains to be determined. The fact that intrathecal CD20 mAb treatment was sufficient to deplete B cells and prevent TLS formation at the DRG suggests a local source of these cells; nonetheless, the origin of the cells populating meningeal TLSs requires further investigation.

A limitation of this work is that intrathecal treatments could also affect other sites in the lumbar region (e.g., inguinal lymph nodes) besides the DRG. However, our previous work did not show an increase in IgG^+^ B cells in draining lymph nodes after PNI^9^. Here, we show a clear association between neuropathic pain and B cell-rich TLSs in the DRG meninges.

Much of the work examining neuroimmune interactions in the DRG has focused on actions of cells within the parenchyma. Recently, dendritic cells and T cells residing in the meninges of the DRG and spinal nerve have been shown to influence activity of sensory neurons and spinal nociceptive neurons ^7, 8^. We further integrate these findings to demonstrate that macrophages, dendritic cells, T cells, and B cells cluster in TLSs in the DRG meninges, organized by germinal center B cells. These structures may be sites whereby autoreactive B cells secrete IgG to activate Fc gamma receptor-expressing cells in the DRG and spinal cord, ultimately leading to neuronal hyperexcitability and pain^9^. Therefore, approaches to disrupt TLSs (e.g., by locally inhibiting B cell activation) could be explored as a powerful potential strategy to alleviate the intractable symptoms of peripheral neuropathic pain.

## ACKNOWLEDGEMENTS

This work was supported by National Institutes of Health grants R01NS126252 (P.M.G.), U19NS130608 (T.J.P.), UC2AR082186 (A.M.M. and M.K.L.), and P30CA016672 (MD Anderson Cancer Center Support Grant, including animal housing and care in the Research Animal Support Facility, Flow Cytometry & Cellular Imaging Core Facility and image analysis in the Advanced Microscopy Core Facility); and by the Assistant Secretary of Defense for Health Affairs endorsed by the Department of Defense, through the Peer Reviewed Medical Research Program under Award Number W81XWH-19-1-0160 and HT9425-24-1-0109 (P.M.G.) (the opinions, interpretations, conclusions, and recommendations contained herein are those of the authors and are not necessarily endorsed by the Department of Defense); the Rita Allen Foundation Award in Pain (to P.M.G.); and, FAPESP: Center For Research in Inflammatory Diseases under grant agreement 2013/08216-2 (to T.M.C.). The murine anti-CD20 monoclonal antibody and IgG2a isotype control antibody were generously gifted by Genentech under a material transfer agreement. We are grateful to the human tissue donors and their families for their gift. We acknowledge the University of Texas at Dallas Genome Core Research facility for their assistance with bulk RNA sequencing and library preparation, and to the University of Texas at Dallas Histology Core Research Facility for providing access to the 10x Xenium instrumentation.

## AUTHOR CONTRIBUTIONS

Conceptualization: T.K.A., C.J.H., P.M.G.

Investigation: T T.K.A., V.K.P., K.F.W., N.T.F., S.M.Z., D.M.R., Y.A.Z., M.J.L., P.S. (mouse in-life procedures); T.K.A., K.F.W., S.M.Z., D.M.R., Y.A.Z. (animal husbandry and colony management); T.K.A. (whole-mount DRG immunostaining, confocal microscopy); V.K.P., T.K.A. (flow cytometry); K.F.W. (ELISAs, tetanus toxoid vaccination); T.K.A. (RT-qPCR); G.V.L.-S., T.M.C., R.M. (mouse scRNA-seq, generation and analysis); J.A.O., A.M.B., J.B.L., T.J.P., L.M.F.J., M.K.L., R.E.M., A.-M.M. (human DRG bulk RNA-seq reanalysis, spatial transcriptomics, and BCR/TCR clonotype analysis); A.V.V., H.F.C. (piglet studies).

Formal analysis: T.K.A. (image analysis); G.V.L.-S., T.M.C., R.M. (mouse scRNA-seq analysis); J.A.O., A.M.B., (human DRG bulk RNA-seq reanalysis, spatial transcriptomics, and BCR/TCR clonotype analysis); all authors (data analysis and interpretation)

Supervision: H.F.C., T.J.P., T.M.C., C.J.H., P.M.G.

Writing – original draft: T.K.A., C.J.H., P.M.G.

Writing – review & editing: all authors

## DATA AVAILABILITY

All data are included in the article and/or supporting information. The mouse scRNA-seq data are available at https://doi.org/10.5281/zenodo.18609123. Raw data for newly generated bulk RNA sequencing of human TLS-containing DRG is available at Gene Expression Omnibus (GEO, accession GSE329721).

## CODE AVAILABILITY

Code for the analysis of mouse scRNA-seq data is available at https://github.com/GabrielLucenaSilva/DRG-meninges-TLS-project.

## REFERENCES

1. Finnerup, N. B., Kuner, R. & Jensen, T. S. Neuropathic Pain: From Mechanisms to Treatment. Physiol Rev 101, 259–301 10.1152/physrev.00045.2019 (2021).

2. McMahon, S. B., La Russa, F. & Bennett, D. L. Crosstalk between the nociceptive and immune systems in host defence and disease. Nat Rev Neurosci 16, 389–402 10.1038/nrn3946 (2015).

3. Ji, R. R., Chamessian, A. & Zhang, Y. Q. Pain regulation by non-neuronal cells and inflammation. Science 354, 572–577 10.1126/science.aaf8924 (2016).

4. Jain, A., Hakim, S. & Woolf, C. J. Immune drivers of physiological and pathological pain. J Exp Med 221, 10.1084/jem.20221687 (2024).

5. Grace, P. M., et al. The Neuroimmunology of Chronic Pain: From Rodents to Humans. J Neurosci 41, 855–865 10.1523/JNEUROSCI.1650-20.2020 (2021).

6. Grace, P. M., Hutchinson, M. R., Maier, S. F. & Watkins, L. R. Pathological pain and the neuroimmune interface. Nat Rev Immunol 14, 217–231 10.1038/nri3621 (2014).

7. Maganin, A. G., et al. Meningeal dendritic cells drive neuropathic pain through elevation of the kynurenine metabolic pathway in mice. J Clin Invest 132, 10.1172/JCI153805 (2022).

8. Du, B., et al. CD4+ alphabeta T cell infiltration into the leptomeninges of lumbar dorsal roots contributes to the transition from acute to chronic mechanical allodynia after adult rat tibial nerve injuries. J Neuroinflammation 15, 81 10.1186/s12974-018-1115-7 (2018).

9. Lacagnina, M. J., et al. B cells drive neuropathic pain-related behaviors in mice through IgG-Fc gamma receptor signaling. Sci Transl Med 16, eadj1277 10.1126/scitranslmed.adj1277 (2024).

10. Fiore, N. T., et al. Reducing IgG accumulation via neonatal Fc receptor (FcRn) blockade relieves neuropathic pain. Brain Behav Immun 125, 371–387 10.1016/j.bbi.2025.01.015 (2025).

11. Fiore, N. T., et al. Autoreactive immunoglobulin G levels and Fc receptor gamma subunit upregulation drive mechanical allodynia after nerve constriction or crush injury. Pain 166, 2804–2817 10.1097/j.pain.0000000000003734 (2025).

12. Pigatto, G. R., et al. GM-CSF instructs vertebral emergency myelopoiesis in neuropathic pain. bioRxiv, 2025.2011.2023.690022 10.1101/2025.11.23.690022 (2025).

13. Sun, B., Ramberger, M., O’Connor, K. C., Bashford-Rogers, R. J. M. & Irani, S. R. The B cell immunobiology that underlies CNS autoantibody-mediated diseases. Nat Rev Neurol 16, 481–492 10.1038/s41582-020-0381-z (2020).

14. Lee, D. S. W., Rojas, O. L. & Gommerman, J. L. B cell depletion therapies in autoimmune disease: advances and mechanistic insights. Nat Rev Drug Discov 20, 179–199 10.1038/s41573-020-00092-2 (2021).

15. Pipi, E., et al. Tertiary Lymphoid Structures: Autoimmunity Goes Local. Front Immunol 9, 1952 10.3389/fimmu.2018.01952 (2018).

16. Drayton, D. L., Liao, S., Mounzer, R. H. & Ruddle, N. H. Lymphoid organ development: from ontogeny to neogenesis. Nat Immunol 7, 344–353 10.1038/ni1330 (2006).

17. Sato, Y., Silina, K., van den Broek, M., Hirahara, K. & Yanagita, M. The roles of tertiary lymphoid structures in chronic diseases. Nat Rev Nephrol 19, 525–537 10.1038/s41581-023-00706-z (2023).

18. Yang, C., Cai, Y. X., Wang, Z. F., Tian, S. F. & Li, Z. Q. Tertiary lymphoid structures in the central nervous system. Trends Mol Med, 10.1016/j.molmed.2024.10.014 (2024).

19. Zhao, L., et al. Tertiary lymphoid structures in diseases: immune mechanisms and therapeutic advances. Signal Transduct Target Ther 9, 225 10.1038/s41392-024-01947-5 (2024).

20. Cohen, M., et al. Meningeal lymphoid structures are activated under acute and chronic spinal cord pathologies. Life Sci Alliance 4, 10.26508/lsa.202000907 (2021).

21. Guillaume, S. M., Beccaria, C. G., Iannacone, M. & Linterman, M. A. Tertiary Lymphoid Structures Across Organs: Context, Composition, and Clinical Levers. Immunol Rev 335, e70063 10.1111/imr.70063 (2025).

22. Meylan, M., et al. Tertiary lymphoid structures generate and propagate anti-tumor antibody-producing plasma cells in renal cell cancer. Immunity 55, 527–541 e525 10.1016/j.immuni.2022.02.001 (2022).

23. Raaphorst, F. M., et al. Coexpression of BMI-1 and EZH2 polycomb group genes in Reed-Sternberg cells of Hodgkin’s disease. Am J Pathol 157, 709–715 10.1016/S0002-9440(10)64583-X (2000).

24. Velichutina, I., et al. EZH2-mediated epigenetic silencing in germinal center B cells contributes to proliferation and lymphomagenesis. Blood 116, 5247–5255 10.1182/blood-2010-04-280149 (2010).

25. Beguelin, W., et al. EZH2 is required for germinal center formation and somatic EZH2 mutations promote lymphoid transformation. Cancer Cell 23, 677–692 10.1016/j.ccr.2013.04.011 (2013).

26. Beguelin, W., et al. EZH2 enables germinal centre formation through epigenetic silencing of CDKN1A and an Rb-E2F1 feedback loop. Nat Commun 8, 877 10.1038/s41467-017-01029-x (2017).

27. Guo, M., et al. EZH2 Represses the B Cell Transcriptional Program and Regulates Antibody-Secreting Cell Metabolism and Antibody Production. J Immunol 200, 1039–1052 10.4049/jimmunol.1701470 (2018).

28. Li, W. W., et al. Autoimmunity contributes to nociceptive sensitization in a mouse model of complex regional pain syndrome. Pain 155, 2377–2389 10.1016/j.pain.2014.09.007 (2014).

29. Dawes, J. M., et al. Immune or Genetic-Mediated Disruption of CASPR2 Causes Pain Hypersensitivity Due to Enhanced Primary Afferent Excitability. Neuron 97, 806–822 e810 10.1016/j.neuron.2018.01.033 (2018).

30. Bersellini Farinotti, A., et al. Cartilage-binding antibodies induce pain through immune complex-mediated activation of neurons. J Exp Med 216, 1904–1924 10.1084/jem.20181657 (2019).

31. Wang, L., et al. Neuronal FcgammaRI mediates acute and chronic joint pain. J Clin Invest 129, 3754–3769 10.1172/JCI128010 (2019).

32. Cuhadar, U., et al. Autoantibodies produce pain in complex regional pain syndrome by sensitizing nociceptors. Pain 160, 2855–2865 10.1097/j.pain.0000000000001662 (2019).

33. Liu, F., et al. Fcgamma Receptor I-Coupled Signaling in Peripheral Nociceptors Mediates Joint Pain in a Rat Model of Rheumatoid Arthritis. Arthritis Rheumatol 72, 1668–1678 10.1002/art.41386 (2020).

34. Goebel, A., et al. Passive transfer of fibromyalgia symptoms from patients to mice. J Clin Invest 131, 10.1172/JCI144201 (2021).

35. Lee, H. J., et al. Sex-Specific B Cell and Anti-Myelin Autoantibody Response After Peripheral Nerve Injury. Front Cell Neurosci 16, 835800 10.3389/fncel.2022.835800 (2022).

36. Guo, T. Z., et al. Pronociceptive autoantibodies in the spinal cord mediate nociceptive sensitization, loss of function, and spontaneous pain in the lumbar disk puncture model of chronic back pain. Pain 164, 421–434 10.1097/j.pain.0000000000002725 (2023).

37. Dieude, M., et al. Extracellular vesicles derived from injured vascular tissue promote the formation of tertiary lymphoid structures in vascular allografts. Am J Transplant 20, 726–738 10.1111/ajt.15707 (2020).

38. Worbs, T., Hammerschmidt, S. I. & Forster, R. Dendritic cell migration in health and disease. Nat Rev Immunol 17, 30–48 10.1038/nri.2016.116 (2017).

39. Rishal, I. & Fainzilber, M. Axon-soma communication in neuronal injury. Nat Rev Neurosci 15, 32–42 10.1038/nrn3609 (2014).

40. Shubayev, V. I. & Myers, R. R. Axonal transport of TNF-alpha in painful neuropathy: distribution of ligand tracer and TNF receptors. J Neuroimmunol 114, 48–56 10.1016/s0165-5728(00)00453-7 (2001).

41. Renthal, W., et al. Transcriptional Reprogramming of Distinct Peripheral Sensory Neuron Subtypes after Axonal Injury. Neuron 108, 128–144 e129 10.1016/j.neuron.2020.07.026 (2020).

42. Pandey, V. K., et al. Peripheral nerve injury reduces macrophage efferocytosis to facilitate neuropathic pain. Proc Natl Acad Sci U S A 123, e2511401122 10.1073/pnas.2511401122 (2026).

43. Tumanov, A. V., Kuprash, D. V., Mach, J. A., Nedospasov, S. A. & Chervonsky, A. V. Lymphotoxin and TNF produced by B cells are dispensable for maintenance of the follicle-associated epithelium but are required for development of lymphoid follicles in the Peyer’s patches. J Immunol 173, 86–91 10.4049/jimmunol.173.1.86 (2004).

44. Uwamizu, A., et al. Autocrine/paracrine lysophosphatidylserine signaling suppresses B cell aggregation and tertiary lymphoid structure formation. iScience 28, 112420 10.1016/j.isci.2025.112420 (2025).

45. Brioschi, S., et al. Heterogeneity of meningeal B cells reveals a lymphopoietic niche at the CNS borders. Science 373, 10.1126/science.abf9277 (2021).

46. Cugurra, A., et al. Skull and vertebral bone marrow are myeloid cell reservoirs for the meninges and CNS parenchyma. Science 373, 10.1126/science.abf7844 (2021).

47. Herisson, F., et al. Direct vascular channels connect skull bone marrow and the brain surface enabling myeloid cell migration. Nat Neurosci 21, 1209–1217 10.1038/s41593-018-0213-2 (2018).

48. Mazzitelli, J. A., et al. Cerebrospinal fluid regulates skull bone marrow niches via direct access through dural channels. Nat Neurosci 25, 555–560 10.1038/s41593-022-01029-1 (2022).

49. Ray, P. R., et al. RNA profiling of human dorsal root ganglia reveals sex differences in mechanisms promoting neuropathic pain. Brain 146, 749–766 10.1093/brain/awac266 (2023).

