## Supplemental Materials for "DRG meningeal tertiary lymphoid structures are regulated by B cells as a pronociceptive locus after peripheral nerve injury"

Tusar K. Acharya *et al.*

**The PDF file includes:**

Materials and Methods
Figs. S1 to S16
Tables S1 to S3
Methods-only references (50-66)

### MATERIALS AND METHODS

#### Animals

All experiments were carried out using male and female mice on a C57BL/6J background, except where indicated. Wild-type mice were obtained from the Jackson Laboratory (C57BL/6J, strain #000664). For experiments with mice lacking mature B cells, muMT mice were bred (B6.129S2-*Ighm*$^{tm1Cgn}$/J, strain #002288, The Jackson Laboratory) with wild-type littermates used as controls. *Aicda*$^{creERT2/+}$::*Ezh2*$^{fl/fl}$ mice were generated by crossing *Aicda*$^{creERT2/+}$ mice (B6.129P2-*Aicda*$^{tm1.1(cre/ERT2)Crey}$/J, The Jackson Laboratory) and *Ezh2*$^{fl/fl}$ mice (B6;129S1-*Ezh2*$^{tm2Sho}$*/J*, strain #022616, The Jackson Laboratory). *Ezh2*$^{fl/fl}$ littermates null for the cre allele were used as control. The animals were administered with tamoxifen (10 mg/kg) prepared in corn oil by oral gavage, for four consecutive days post PNI surgery. Upon arrival at the pathogen-free, AAALAC accredited MD Anderson vivarium, mice were given at least 1 week to acclimate to environmental conditions before any manipulations were performed. Mice of all strains were housed 2-5 per cage in individually ventilated cages (14 in x 7 in x 7 in, Tecniplast) with *ad libitum* access to food (Purina PicoLab Rodent Diet 5053) and drinking water. The housing conditions of each cage included absorbable corncob bedding and cotton fiber nesting material, and mice were transferred to clean cages once every 2 weeks. The colony was maintained in temperature- and humidity-controlled rooms on a 12 h light-dark cycle (lights on from 7:00 to 19:00 hours), and all procedures were performed during the light cycle. Subjects were ordered from vendors or bred in-house at the University of Texas MD Anderson Cancer Center, and mice were at least 8 weeks old at the start of experimental manipulations. Mice were randomly and prospectively assigned to experimental groups using an online random number generator (https://www.randomlists.com/team-generator). Investigators were blinded to experimental group assignments during data collection and analysis. All animal experiments and procedures were approved by the Institutional Animal Care and Use Committee of the University of Texas MD Anderson Cancer Center.

Commercial crossbred porcine piglets (1 day old at arrival; approximately 1 kg body weight) were obtained and housed at Midwest Veterinary Services–KS (Manhattan, KS, USA; Protocol No. MVS 23001). All procedures were approved by the MVS-KS Institutional Animal Care and Use Committee (IACUC) and conducted in accordance with the *Guide for the Care and Use of Agricultural Animals in Research and Teaching (Ag Guide)*. A total of 40 piglets (20 males and 20 females) were acclimated for 2 days prior to study initiation, and cross-fostering was permitted to achieve the required sex distribution. Within each litter, the five heaviest healthy pigs per sex were selected for study enrollment, while non-enrolled animals were humanely euthanized according to AVMA-approved methods and institutional standard operating procedures. Animals were housed with sows in stainless steel farrowing crates equipped with automatic watering systems, self-feeding troughs, heat lamps, heat mats, and rubber mats, with housing conditions meeting or exceeding the standards outlined in the Guide for the Care and Use of Agricultural Animals in Agricultural Use and Research and Teaching 4th Edition. Environmentally controlled conditions, including a 14 h light/10 h dark photoperiod, temperature-controlled housing, *ad libitum* access to pelleted diet formulated for pigs and water were maintained throughout the study. Animals underwent daily health observations by trained personnel.

#### PNI surgeries

PNI was induced by chronic constriction injury (first model of PNI) for all experiments except for the results reported in transcriptomic analysis (Fig. 5 and S9), where spared nerve injury was used. Chronic constriction injury of the sciatic nerve[50], was performed as we have described[9]. Under isoflurane anesthesia (2-3% in oxygen), the midthigh level of the left leg was shaved, cleansed with povidone-iodine and 70% ethanol, and the sciatic nerve exposed through blunt dissection of the biceps femoris muscle. Three ligatures (5-0 chromic gut; Ethicon) were loosely tied around the sciatic nerve. For sham surgeries, the sciatic nerve was isolated, but no chromic gut sutures were applied. Spared nerve injury (second model of PNI)[51, 52] was performed as we have described[53]. Under inhaled isoflurane aesthesia, the tibial and common peroneal nerves were isolated, tightly ligated with 6-0 silk (707G, Ethicon, Somerville, USA), and transected immediately distal to the ligation. The sural nerve was left intact. The muscle layer was sutured closed with silk, and the skin was closed with wound clips. For sham surgery, the nerves were exposed, but not ligated or transected. At the completion of surgery, mice were returned to their home cage and monitored postoperatively until fully ambulatory. For experiments in which naive subjects were used, mice were left undisturbed in their home cages.

For the tail docking surgery, the piglets underwent tail docking or sham docking at 5 days old. Tail docking was conducted with one person holding the piglet and a second person removing half of the piglet's tail with side pliers. Sham docking was conducted by mimicking the tail docking procedure with fingers instead of pliers. After surgery, the animals were monitored under supervision.

**Immunostaining and analysis of whole mount DRG and spine sections**

Either 14 or 28 days after PNI/sham surgery, mice were euthanized using a lethal injection of pentobarbital-phenytoin solution intraperitoneal (i.p.) and transcardially perfused with 10 mL of ice-cold PBS, followed by 4% paraformaldehyde (PFA). The ipsilateral and contralateral L4-L5 DRGs with intact meninges were carefully harvested and post-fixed in 4% PFA for 24 hours. The tissues were then transferred to a 30% sucrose solution and stored at 4°C until further processing. Wholemount immunostaining was performed as a multistep process, with all procedures conducted under floating conditions at 4°C to preserve tissue integrity. The DRGs were first incubated in 0.1% Triton X-100 in PBS (cat# T9284, Millipore Sigma, USA) for 24 hours to permeabilize the tissues. The tissues were blocked with 5% donkey (cat# ab7475, Abcam, USA) or goat (cat# 50062Z, Invitrogen, USA) serum for 4 hours, and then incubated with primary antibodies for 48 hours (see Table S1 for vendor and dilution information). The tissues were washed multiple times over a 12-24-hour period with 0.1% Triton X-100, and then incubated with secondary antibodies for 12 hours see Table S1 for vendor and dilution information), followed by another 12-24 hours of washing with 0.1% Triton X-100. To label nuclei, the tissues were incubated with DAPI for 15 minutes. After staining, the tissues were mounted using FluorSave reagent (cat#345789, Millipore Sigma, USA) and allowed to dry at 4°C for 2 days. The specificity of IgG immunolabeling was confirmed using antibody controls (Fig. S14a).

For DRG collection from piglets, animals were euthanized at 14 or 42 days post-surgery by captive bolt followed by exsanguination. DRGs were harvested, drop-perfused in 10% formalin, transferred to 30% sucrose solution and stored at 4°C until further processing. Wholemount immunostaining was performed as described above.

For the staining of spine sections, mice were euthanized 14 days after PNI surgery using a lethal injection of pentobarbital-phenytoin solution intraperitoneal (i.p.), and then transcardially perfused with 10 mL of ice-cold PBS, followed by 4% PFA. Spinal columns were post-fixed in 4% PFA for 48 hours at

4° C, then washed with 1x PBS for 24 hours at 4° C. Decalcification was performed using 10% EDTA (cat. E9884, Millipore Sigma, USA) at pH 7.4 for 15 days with the solution changed every 3 days. Upon complete decalcification, tissues were stored in PBS containing 0.02% sodium azide at 4° C until further processing. Decalcified spines were cryo-sectioned at a thickness of 30 µm. Immunohistochemistry of spine sections was performed by using antibodies against B220, CD45 and podoplanin.

Images were captured with a Nikon Eclipse Ti2 confocal microscope using 10x and 40x objective with a pinhole of 1AU and 1024×1024 pixel size at 1.5 µm stack size. The Z stack images were merged by using maximum projection function by ImageJ. For enhanced visualization, pseudo coloring has been applied to the images, and some digitally zoomed images have been rotated for better clarity and representation. For quantification, the images were captured at 40x magnification with an 8×8 tiling format at 1024×1024-pixel resolution maintaining 1.5µm Z stack interval. The image processing and quantification were performed using ImageJ and Imaris software (Version 8.4.2). $CD45^+$ immune cell clusters were quantified from 40x tiled images using an automated surface-based workflow in Imaris (Oxford Instruments), adapted from previously described methods[54]. Surface-based segmentation was performed using an identical absolute intensity threshold and a smoothing factor of 0.621 µm to minimize background noise while preserving true signal and generating individual segmented objects. Segmented objects were automatically classified as small aggregates, medium clusters, or large clusters according to increasing surface area statistics using the built-in one-dimensional (1D) classification module in Imaris without manual assignment of size thresholds or adjustment of class boundaries. Small aggregates, comprising the smallest segmented objects, were excluded from downstream analyses as they could not be reliably distinguished from single cells, whereas medium and large clusters were retained as $CD45^+$ immune cell clusters. The median and observed range of surface area, volume, sphericity, and ellipticity (oblate/prolate) for medium and large clusters are presented in Table S2; these values describe the characteristics of the automatically classified objects and were not used as classification criteria. Sphericity and ellipticity (oblate/prolate) values range from 0-1, where higher values indicate greater sphericity or ellipticity; prolate ellipticity corresponds to elongated clusters, whereas the oblate ellipticity corresponds to flattened clusters. All segmentation, thresholding, and classification parameters were established before analysis and applied identically to all samples using the same automated analysis pipeline without sample-specific optimization.

For the quantification of immune cell numbers, ~6-7 regions of interest (ROIs) were manually selected from each DRGs based on areas of prominent immune infiltration. The cell numbers were quantified using ImageJ software (v.1.54p) within a fixed ROI area of 318.2 × 318.2 µm (0.10125 $mm^2$), and the data are presented as average cells/$mm^2$. Because individual macrophages and dendritic cells could not be resolved based on morphology, the intensity of $F4/80^+$ and $CD11c^+$ staining was measured. The mean gray intensity for each ROIs was quantified using ImageJ software and the average mean gray intensity was plotted. For podoplanin and Lyve1, integrated density was measured across the entire DRG tissue to quantify total intensity.

**Flow cytometry**

Mice were euthanized using a lethal injection of pentobarbital-phenytoin solution intraperitoneal (i.p.). Single cell suspensions were prepared from the pooled L3-5 DRGs with attached meninges (three mice per sample). A total of eight pooled samples per experimental group were analyzed. Tissues were enzymatically dissociated by using digestion medium containing DMEM, HEPES (10 mM), BSA (5mg/mL), collagenase IV (2mg/mL), DNase I (100 µg/mL) at 37 °C for 40 min with continuous agitation (≤ 600 rpm). Following enzymatic digestion, tissues were gently triturated 10–15 times using a 1-mL pipette tip to obtain a single-cell suspension. The resulting suspension was passed through 70 µm cell strainer and washed using 10-fold excess volume of FACS buffer (PBS with 5% FBS and 1mM EDTA) by centrifugation at 300 x g for 10 min at 4 °C. Cell pellets were resuspended in FACS buffer for subsequent staining.

For surface receptor staining, cells were stained with fluorophore-conjugated antibodies (Table S1) for 30 min at 4 °C. Following surface staining, cells were washed with FACS buffer, followed by PBS and then incubated with viability dye (Table S1) for 30 min. Cells were subsequently washed with FACS buffer and fixed using 2% formaldehyde for 30 min at room temperature. For intracellular labelling, the fixed cells were permeabilized by using intracellular staining perm/wash buffer (cat# 421002, Bio Legend) for 10 min in ice, followed by incubation with intracellular antibodies (Table S1) overnight at 4 °C. The next day, the cells were washed with perm/wash Buffer, resuspended in FACS buffer, and acquired on a Gallios flow cytometer (Beckman Coulter, Brea, CA, USA) or a CytoFLEX LX flow cytometer (Beckman Coulter, Brea, CA, USA) at The University of Texas MD Anderson Cancer Center Flow Cytometry & Cellular Imaging Core Facility. Flow cytometry data were analyzed using Kaluza 2.2 software (Beckman Coulter) or FlowJo v10 software (BD Biosciences). Cells were gated based on forward scatter (FSC) and side scatter (SSC) to exclude debris followed by singlet discrimination to remove cell aggregates and live cell gating to exclude dead cells. $CD45^+$ cells were identified by using the corresponding fluorescent minus one (FMO) control. All subsequent immune cell populations were analyzed within $CD45^+$ population using their respective FMO controls (S15).

**Single-cell RNA sequencing (scRNA-seq) data analyses of mouse tissues**

The scRNA-seq data from DRG meningeal leukocytes ($CD45^+$ cells) has been acquired and analyzed previously[12]. Raw data are available at https://doi.org/10.5281/zenodo.18609123. Herein, cells assigned to the B cell lineage were extracted from the integrated dataset and analyzed independently. Unlike the full dataset, which was normalized with Log Normalize and imputed with ALRA, the B cell subset was re-normalized using SC Transform (3,000 variable features), regressing out mitochondrial content. PCA was performed on the SCT-corrected data, and the first 30 components were retained. A shared nearest-neighbor graph was built from these components (k = 10 nearest neighbors), and cells were clustered with the Louvain algorithm at a lower resolution (0.5) than the full atlas, to resolve biologically coherent B cell states without over-fragmentation. Subclusters were visualized by t-distributed stochastic neighbor embedding (t-SNE; 30 components, 30 nearest neighbors). Marker genes for each subcluster were identified with FindAllMarkers using the Wilcoxon rank-sum test (positive markers only; min.pct = 0.25; log2 fold-change threshold ≥ 0.25). Subclusters were manually annotated into eight B cell states: Immature, Naive Follicular, Mature Follicular, APC-like, Pre-GC, GC, Activated, and Mature B cells, based on curated marker-gene modules. All plots were generated using the R package ggplot2 [55]. Color palettes were from the R packages Pals [56].

Gene Ontology Biological Process (GO BP) enrichment analysis was performed on differentially expressed genes with the GO Biological Process 2023 database. For each cluster, the top 100 significantly enriched GO BP terms ranked by *P* value were selected for pathway network visualization. A pathway similarity network was generated in Cytoscape v3.10.4[57] using custom node and edge tables, where nodes represented GO BP pathways and edges represented pathway similarity based on shared genes (Jaccard similarity coefficient). The network was visualized using the EnrichmentMap and AutoAnnotate app collection[58]. Pathways were clustered using the Markov Cluster Algorithm (MCL), and cluster annotations were automatically generated using AutoAnnotate with Node Name as the label source. Network layouts were optimized to minimize cluster overlap, and node size was scaled according to the number of genes associated with each pathway.

**RNA extraction and bulk RNA sequencing in human tissues**

A lumbar disc degeneration-associated human DRG (spinal level L3/L4) was embedded in optimal cutting temperature medium (Sakura). Three 100 µm sections were taken and stored in a 1.5 mL tube at -80°C until RNA extraction. QIAzol (1 mL, Qiagen #79306) was added to the tube containing tissue sections and the contents were transferred to a bead-based tissue homogenization tube (Precellys) and shaken for 1 min at 4°C. The lysed tissue was transferred to a new tube, and 200 µL chloroform was added. The tube was shaken for 15 sec and left to incubate for 2 min at RT. The tube was centrifuged at 12,000 g for 10 min at 4°C. The upper aqueous phase was collected, mixed with the same volume of 70% ethanol, and loaded on a column for RNA extraction using the RNeasy mini kit (Qiagen #74104). At this and future steps, the column was centrifuged at 8000 g for 15 sec at 4°C and the flow-through was discarded. Then, 350 µL RW1 buffer was added, the column was centrifuged, then 80 µL of DNase in RDD buffer was added and left to incubate for 15 min at RT. The sample was washed with 350 µL RW1 buffer, then twice washed with 500 µL RPE buffer. The column was then dried by centrifuging at 16,000 g for 1 min at RT. RNA was eluted in 20 µL RNase-free water by incubating for 1 min and centrifuging into a clean collection tube. RNA quality was verified on a NanoDrop 2000.

Library preparation was performed by the University of Texas at Dallas Genome Core Research Facility using the Illumina Stranded Total RNA Prep kit per the manufacturer's protocol. Sequencing was performed with a read length of 50 bp paired end on an Illumina NextSeq 2000 at the University of Texas at Dallas Genome Core Research Facility. Quality control of the raw FASTQ file was assessed using the Phred score, duplication level, and per-base sequence with FastQC v0.12.1 built from conda 4.14.0.

**Clonotype analyses**

Mature tertiary lymphoid structures contain germinal centers. Using bulk RNA sequencing of human DRG, we can extract relative clonotype numbers by aligning reads to B- and T- cell receptors where present. Raw fastq.gz files were used for receptor immune profiling with MiXCR v4.7.0 [59] (RepSeq.IO v2.5.0, MiLib v3.5.0, repseqio.v5.1), built from conda 24.11.1. For the lumbar disc degeneration-associated human DRG reported in this manuscript, default settings for paired end bulk RNA-seq alignment were used (`mixcr analyse rnaseq`).

Additionally, previously sequenced human DRG samples were used to broaden this to a larger neuropathic pain cohort[49]. A subset of samples (n = 10) were only available as filtered/trimmed bam files which could not be converted to fastq in totality. These samples were thus not processed. Samples (50 bp or 75 bp) were split by read length, due to alignment efficiently, and were analysed with mixcr using default settings for single-end bulk RNA-seq alignment. Only samples considered "neuronal" and with pain scores in the original paper were considered. Here, only samples considered "neuronal" and with pain scores in the original paper were considered. For 75 bp samples, No Pain = 8, NeuP = 14 were processed. For 50 bp samples, No Pain = 5, NeuP = 13.

**Gene set enrichment analyses (GSEA)**

We have previously used single sample gene set enrichment analysis (ssGSEA) to highlight B cell enrichment in human DRG[9]. Here, we exploited additional gene sets to probe for tertiary lymphoid structure signatures in the same cohort – neuropathic pain in human DRG [49]. Gene symbols were converted to ensembl IDs via the `biomaRt::getBM` from https://sep2025.archive.ensembl.org.

TLS_Meylan2022 gene set was used as published[22]. This gene set was first generated in the context of mature TLS in renal cell cancer and has since been validated across additional cancer types[60]. An in-house gene set (TLS_ curated gene list) was also generated as a list of markers across cell types relevant to TLS, which are hypothesized to be enriched in DRG with TLS (Table S3). GSEA was performed against ranked quantile normalized transcripts per million (qnTPM) using the clusterProfiler package in R (v4.18.4)[61]. Positive enrichment was examined per sample (nPermSimple = 10000), as the normalized enrichment score (NES) from ClusterProfiler::GSEA. One donor was excluded for an undocumented pain state, resulting in 17 non-pain DRG and 33 pain DRG.

**Spatial transcriptomics**

10x Xenium imaging-based spatial transcriptomics was performed on a lumbar disc degeneration-associated human DRG using a custom gene panel of 472 genes that has been previously used[62]. One 10 μm section from the embedded DRG was passed on to the 10x Xenium sample preparation and data acquisition workflow, which was performed with cell segmentation staining according to the supplier's standard protocols (CG000581RevD, CG000749RevA, CG000584RevF) and a probe hybridization time of 17 hours. Cell segmentation for neurons was performed manually on staining images in QuPath v0.5.1 and then passed through Xenium Ranger for resegmentation, using nuclei expansion for remaining non-neuronal cells. Representative images of histology and transcript localization were generated in Xenium Explorer v4.1.1.

Resegmented Xenium data were loaded into Seurat v5.4.0 in R v4.4.1. Cells with zero counts were removed and the SCTransform function was applied with default parameters. Louvain clustering was performed at a resolution of 0.3 to generate 20 clusters, which were annotated according to differentially expressed marker genes for broad cell types with reference made to a previously generated human dorsal root ganglion single-nuclei RNA sequencing atlas[63]. Spatial visualizations of annotated cell centroids and associated marker genes were generated using Seurat's ImageDimPlot function.

**Depletion of B cells with CD20 mAb**

CD20 monoclonal antibodies (mouse IgG2a, clone 5D2) and isotype control antibodies were generously provided by Genentech. Mice were administered either monoclonal anti-CD20 Ab or isotype control (10 μg in 5 μL volume prepared in sterile 0.9% saline) through intrathecal injection at the L4-L5 spinal level under isoflurane anesthesia, as described[9, 64]. The dose was selected from preliminary dose response studies. The injections were performed immediately prior to PNI or sham surgeries.

**Intrathecal transfer of B cells**

Splenic B cells were purified from wild-type donor mice by magnetic-activated cell sorting using a B cell isolation kit (cat#130-090-862, Miltenyi Biotec, USA) as we have previously described [9]. muMT mice received 2 x $10^5$ B cells in 5 μL of PBS (or equivolume PBS control) by intrathecal injection at the L4-L5 spinal level. The adoptive transfer of B cells and PBS were carried out immediately before the PNI and sham surgeries based on our previous report[9].

**Behavioral test for mechanical allodynia**

Mice were habituated to the testing apparatus and the experimenter prior to experimentation. The experimenter remained blinded to the group assignment of each animal during the testing period. Von Frey testing of punctate allodynia was performed as previously described[53, 64, 65]. The up-down method was performed and 50% withdrawal thresholds were calculated as previously described[66].

**Tetanus toxoid vaccination**

Tetanus toxoid vaccine (ANTIGEN-1002, Charles River, USA) was administered via intramuscular (i.m.) injection into each hind leg (10 μg in 100 μL per injection) 14 days prior to PNI surgery/CD20 mAb injections. Tetanus-toxoid booster injections were administered 7 days after PNI surgery/CD20 mAb injections (10 μg in 5 μL volume). At 21 days after PNI surgery/CD20 mAb injection, mice received a lethal i.p. injection of pentobarbital-phenytoin solution and blood was collected via cardiac puncture.

Blood was centrifuged at 3200 x g at room temperature for 10 minutes, serum was collected and stored at -80°C.

### ELISA

Serum concentration of anti-tetanus toxoid IgG was assessed using a tetanus toxoid IgG ELISA (930-130-TMG, Alpha diagnostics, TX, USA). Anti-tetanus toxoid IgG ELISA was performed following manufacturer's instructions, with serum diluted 1:30,000 and 100 µL of diluted serum loaded per sample. All samples and standards were run in duplicate, and tetanus toxoid IgG concentration was quantified based on the standard curve. Anti-tetanus toxoid IgG expression was normalized to units per µL serum sample.

### RT-qPCR

B cells were isolated from spleens using B cell isolation kit (cat#130-090-862, Miltenyi Biotec, USA). The total RNA from the B cells were extracted by using RNA mini kit (cat# 12183018A, Invitrogen, USA) according to the manufacturer's instructions. The cDNA was synthesized from 1 µg of total RNA using SuperScript IV VILO Master Mix (cat# 11756050, Invitrogen, USA). Quantitative real time PCR (qPCR) was performed using SYBR Green Supermix (cat#1725274, BioRad, USA) on a BioRad CFX Connect Real-Time PCR system to quantify *Ezh2* transcript levels. The following primers were used: Forward 5' -GGCCTCATAGTGACAGGTCTTAAAA; Reverse 5'- CGCGTCTCCGGATGTACAG. Expression levels were normalized to housekeeping gene *Gapdh*, and relative expression was calculated using the ΔΔCt values (Fig. S16).

### Statistical analysis

Data are presented as group mean ± SEM with individual subjects presented as symbols whenever feasible, unless otherwise indicated. Statistical analyses were conducted in GraphPad Prism (v10). The Shapiro–Wilk test was used to assess normality. The numbers of $CD45^+$ cell clusters were analyzed by unpaired, two-tailed Student's t tests or two-way ANOVA with Tukey's multiple comparison test. Cell numbers or integrated densities were analyzed by unpaired, two-tailed Student's t test or Tukey's multiple comparisons test. Von Frey data were analyzed by repeated measures two-way ANOVA with Sîdak's post hoc tests. ELISA and PCR data were analyzed by unpaired, two-tailed Student's t tests. Differences between groups were considered statistically significant when $p < 0.05$. Exact p values are presented where possible.

### METHODS ONLY REFERENCES


50. Bennett, G. J. & Xie, Y. K. A peripheral mononeuropathy in rat that produces disorders of pain sensation like those seen in man. *Pain* **33**, 87-107 https://doi.org/10.1016/0304-3959(88)90209-6 (1988).
51. Decosterd, I. & Woolf, C. J. Spared nerve injury: an animal model of persistent peripheral neuropathic pain. *Pain* **87**, 149-158 https://doi.org/10.1016/S0304-3959(00)00276-1 (2000).
52. Shields, S. D., Eckert, W. A. & Basbaum, A. I. Spared nerve injury model of neuropathic pain in the mouse: A behavioral and anatomic analysis. *Journal of Pain* **4**, 465-470 https://doi.org/10.1067/S1526-5900(03)00781-8 (2003).
53. Avery, T. D.*, et al.* Site-specific drug release of monomethyl fumarate to treat oxidative stress disorders. *Nat Biotechnol* **43**, 1624-1627 https://doi.org/10.1038/s41587-024-02460-4 (2025).

54. Boulat, V., *et al.* Protocol for rapid 5-plex 3D imaging and single-cell analysis of immune responses in whole murine lymph nodes. *STAR Protoc* **6**, 104059 https://doi.org/10.1016/j.xpro.2025.104059 (2025).
55. Wickham, H. *ggplot2: Elegant Graphics for Data Analysis*, 2 edn. (Springer Cham, 2016).
56. Wright, K. pals: A comprehensive collection of color palettes, colormaps, and tools to evaluate them. R package version 1.10.) (2025).
57. Shannon, P., *et al.* Cytoscape: a software environment for integrated models of biomolecular interaction networks. *Genome Res* **13**, 2498-2504 https://doi.org/10.1101/gr.1239303 (2003).
58. Merico, D., Isserlin, R., Stueker, O., Emili, A. & Bader, G. D. Enrichment map: a network-based method for gene-set enrichment visualization and interpretation. *PLoS One* **5**, e13984 https://doi.org/10.1371/journal.pone.0013984 (2010).
59. Bolotin, D. A., *et al.* MiXCR: software for comprehensive adaptive immunity profiling. *Nat Methods* **12**, 380-381 https://doi.org/10.1038/nmeth.3364 (2015).
60. Li, H., *et al.* Mature tertiary lymphoid structures evoke intra-tumoral T and B cell responses via progenitor exhausted CD4(+) T cells in head and neck cancer. *Nat Commun* **16**, 4228 https://doi.org/10.1038/s41467-025-59341-w (2025).
61. Wu, T., *et al.* clusterProfiler 4.0: A universal enrichment tool for interpreting omics data. *Innovation (Camb)* **2**, 100141 https://doi.org/10.1016/j.xinn.2021.100141 (2021).
62. Sankaranarayanan, I., *et al.* Progressive neurodegeneration in human dorsal root ganglion from diabetes to painful neuropathy. *bioRxiv*, https://doi.org/10.64898/2026.01.16.700028 (2026).
63. Bhuiyan, S. A., *et al.* Harmonized cross-species cell atlases of trigeminal and dorsal root ganglia. *Sci Adv* **10**, eadj9173 https://doi.org/10.1126/sciadv.adj9173 (2024).
64. Zhang, J., *et al.* HDAC6 Inhibition Reverses Cisplatin-Induced Mechanical Hypersensitivity via Tonic Delta Opioid Receptor Signaling. *J Neurosci* **42**, 7862-7874 https://doi.org/10.1523/JNEUROSCI.1182-22.2022 (2022).
65. Li, J., *et al.* Oral Dimethyl Fumarate Reduces Peripheral Neuropathic Pain in Rodents via NFE2L2 Antioxidant Signaling. *Anesthesiology* **132**, 343-356 https://doi.org/10.1097/aln.0000000000003077 (2020).
66. Chaplan, S. R., Bach, F. W., Pogrel, J. W., Chung, J. M. & Yaksh, T. L. Quantitative assessment of tactile allodynia in the rat paw. *J Neurosci Methods* **53**, 55-63 https://doi.org/10.1016/0165-0270(94)90144-9 (1994).

### SUPPLEMENTARY FIGURES

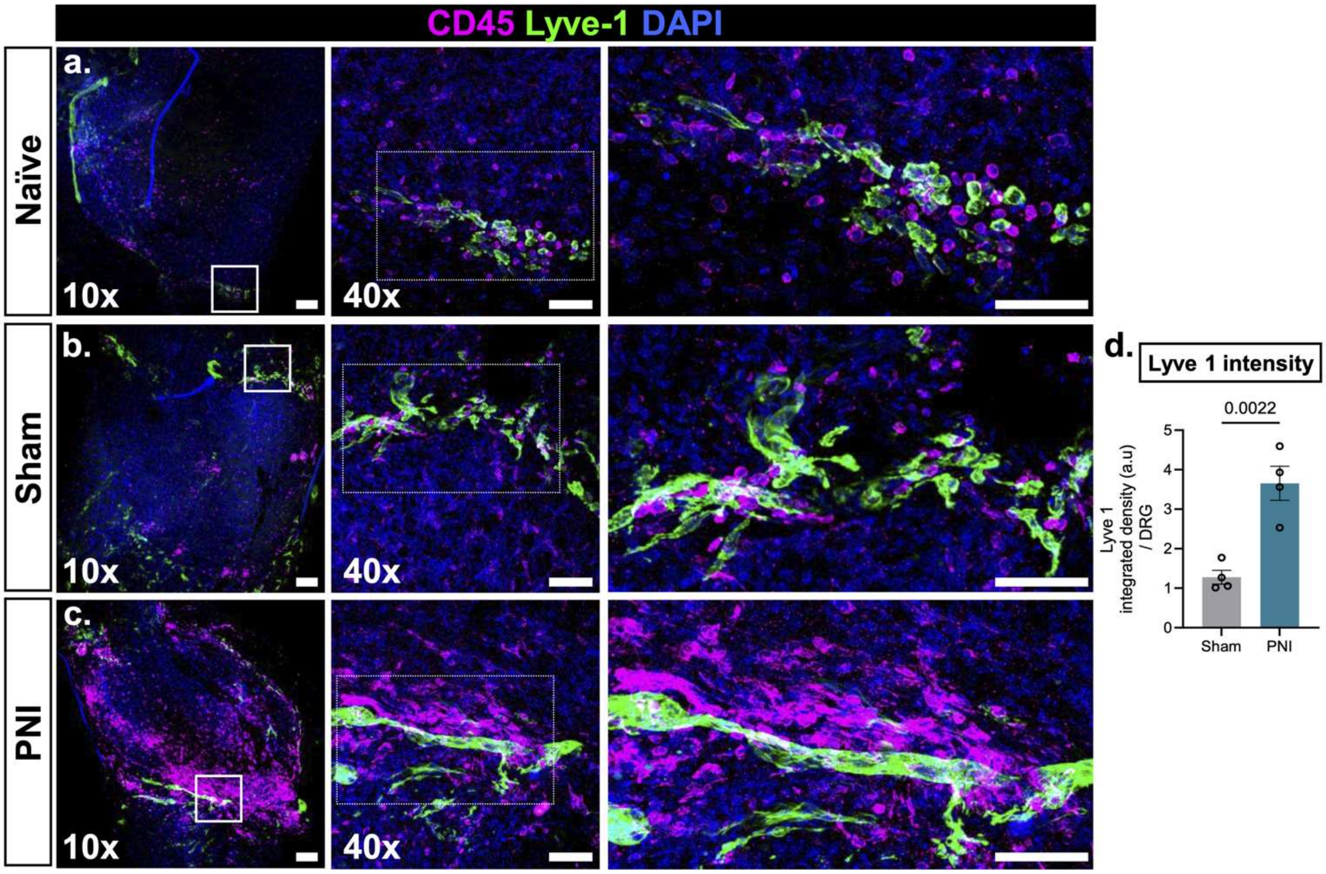


**Figure S1. PNI induces $CD45^+$ cell clusters in the DRG meninges.**
**(a-c)** Representative 10x and 40x (inset) images of whole mount L4-L5 DRGs collected from male mice after 14days of PNI, sham surgery or naïve (scale bar = 100µm). The enlarged images were digitally cropped from the original 40x images (some are rotated, scale bar = 50 µm). DRGs were stained for a pan leukocyte marker (CD45), lymphatic endothelial marker to define the meningeal surface (Lyve-1), and a nuclear marker (DAPI). **(a)** from a naïve mouse **(b)** ipsilateral to sham **(c)** ipsilateral to PNI. **(d)** Integrated density of Lyve-1. Data analyzed by unpaired, two-tailed Student's t tests. N = 4 per group.

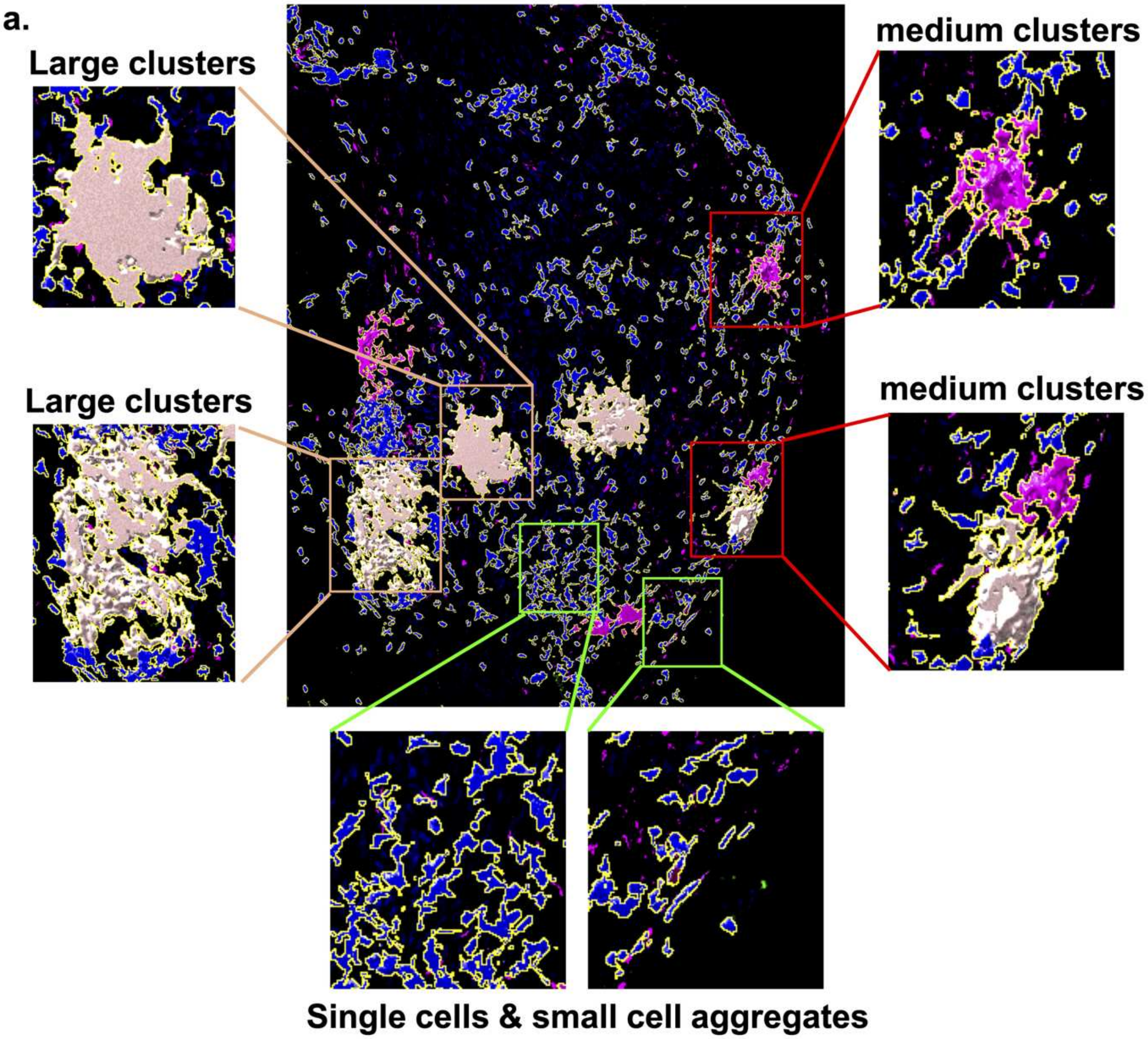


**Figure S2. Surface view (3D) of the DRG surfaces and clusters.**
(**a**) Representative 3D surface renderings of $CD45^{+}$ cell clusters in the DRGs collected 14 days after PNI, illustrating single cells and small aggregates, medium and large clusters.

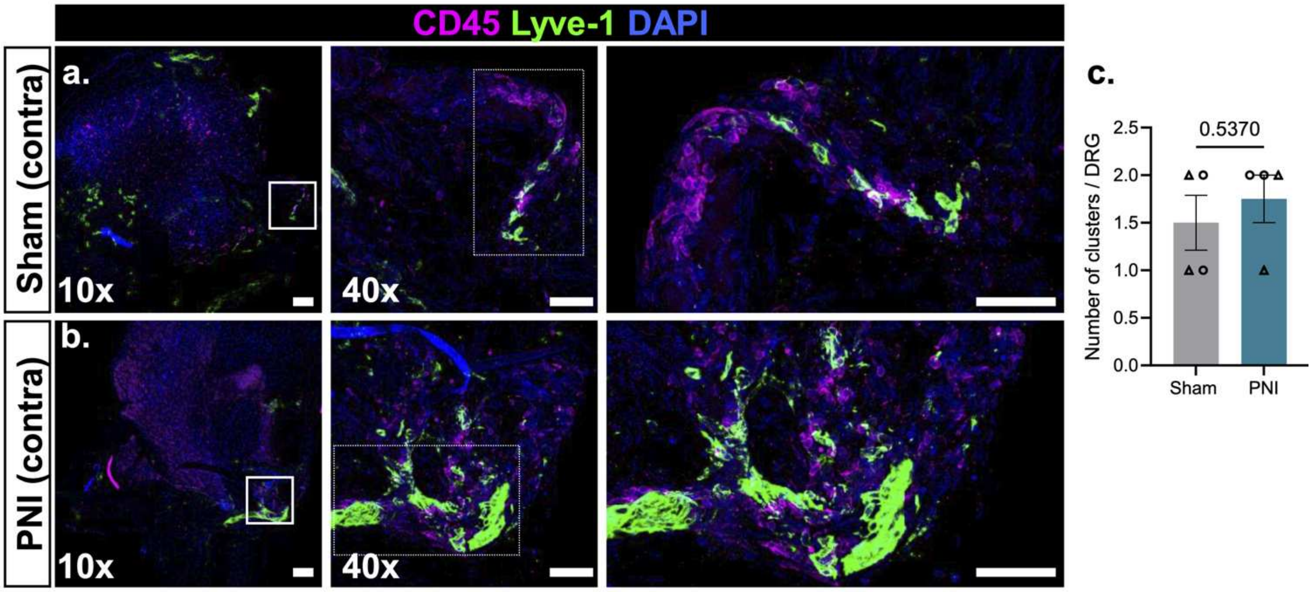


**Figure S3. CD45⁺ cell staining at the contralateral DRGs.**

(**a-b**) Representative 10x and 40x (inset) images of whole mount contralateral L4-L5 DRGs collected from male mice 14 days after (**a**) sham surgery or (**b**) PNI (scale bar = 100 µm). The enlarged images were digitally cropped from the original 40x images (some are rotated, scale bar = 50 µm). DRGs were stained for a pan-leukocyte marker (CD45), lymphatic endothelial marker to define the meningeal surface (Lyve-1), and a nuclear marker (DAPI). **(c)** Number of total clusters around the male and female contralateral DRGs were quantified. N = 4 per group (both sexes included). Data analyzed by unpaired, two-tailed Student's t tests.

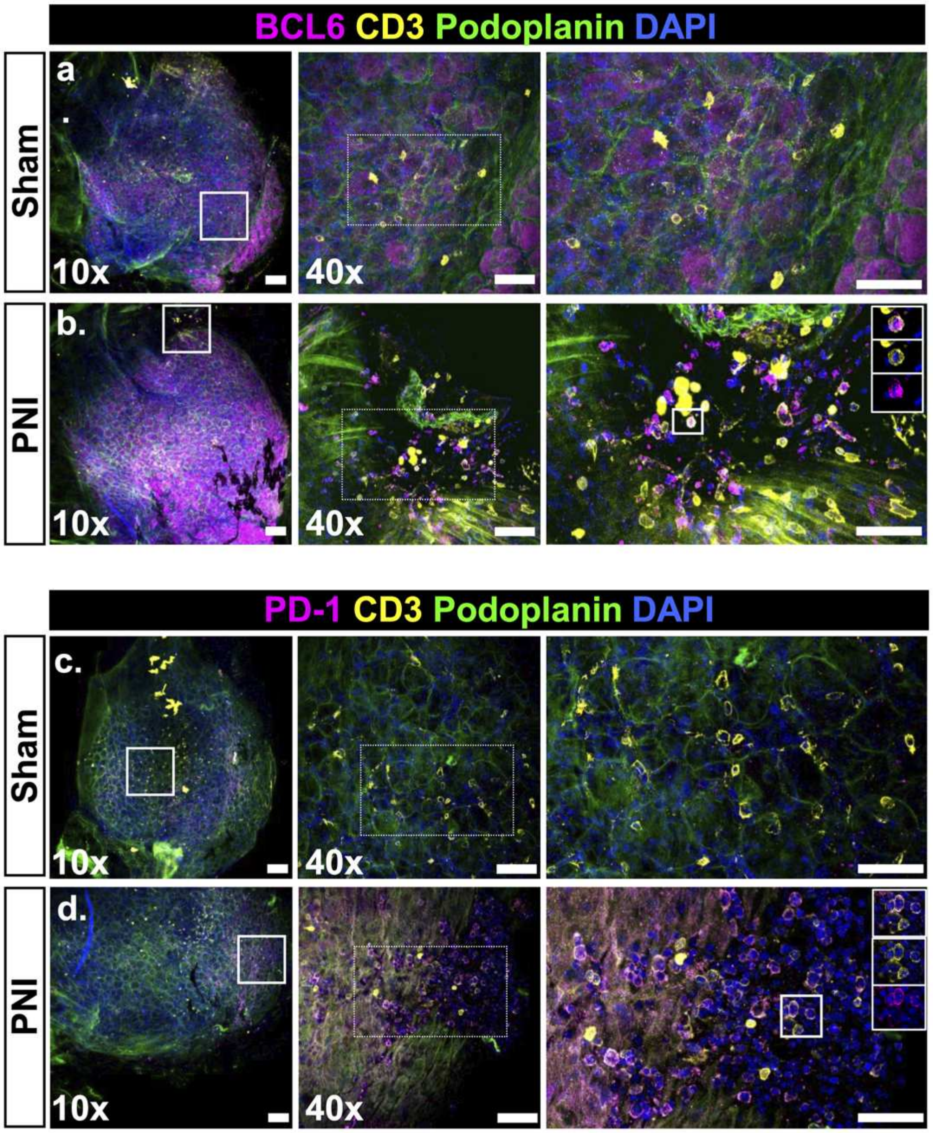


**Figure S4. PNI induces expansion of T follicular helper (Tfh) cells in the DRG meninges.** Representative 10x images with corresponding 40x inset images of whole-mount ipsilateral L4–L5 DRGs with attached meninges collected from male mice 14 days after sham or PNI surgery (scale bar = 100 µm). Enlarged images were digitally cropped from the original 40x images, and some were rotated for presentation (scale bar = 50 µm). **(a-b)** Representative immunofluorescence images showing staining for CD3 (T cells), BCL6 (Tfh cell marker), podoplanin (lymphatic endothelial cells), and DAPI (nuclei). **(c-d)** Representative immunofluorescence images showing staining for CD3 (T cells), PD-1 (Tfh cell marker), podoplanin (lymphatic endothelial cells), and DAPI (nuclei).

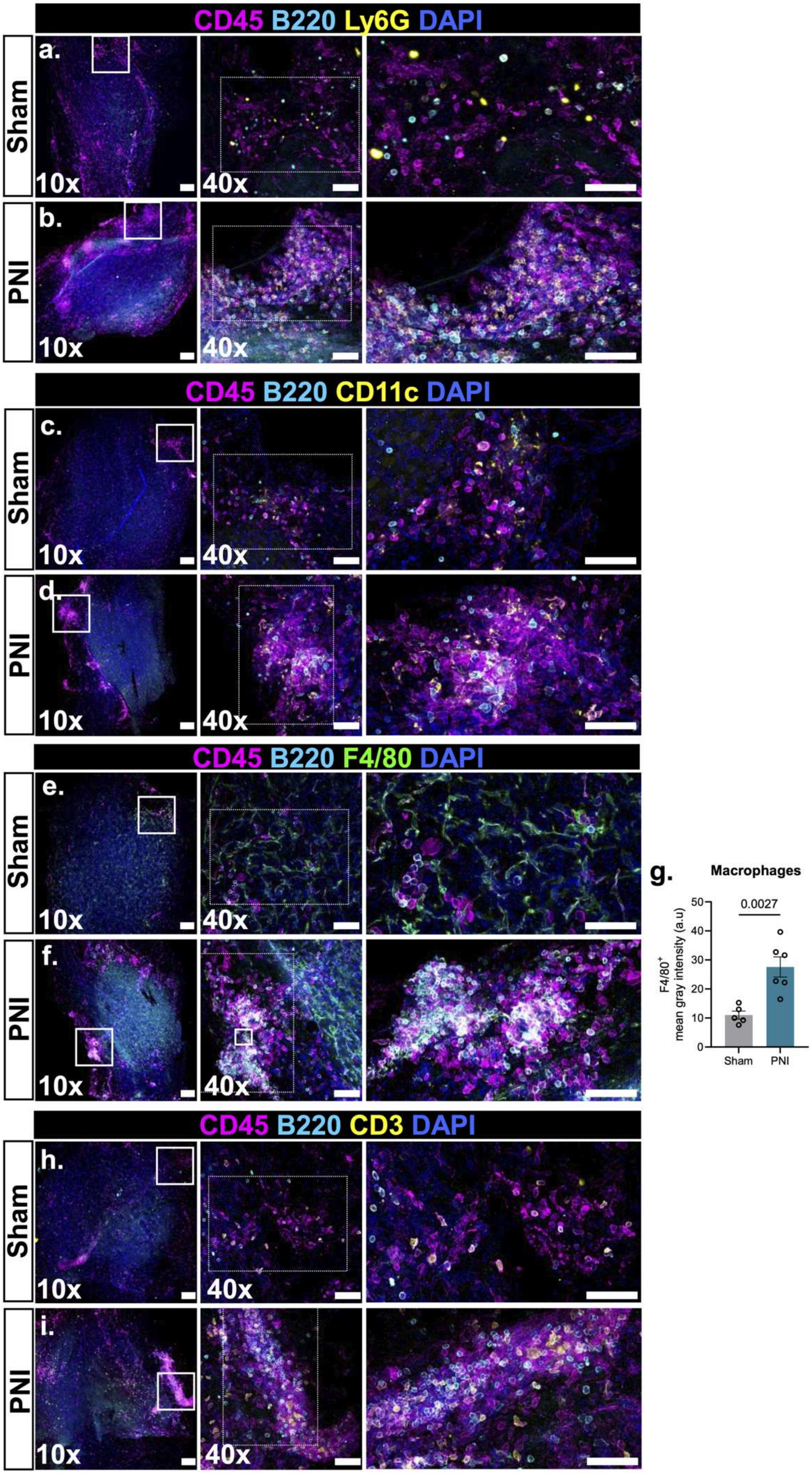
CD45 B220 Ly6G DAPI
a.
Sham
10x
40x
b.
PNI
10x
40x
CD45 B220 CD11c DAPI
c.
Sham
10x
40x
d.
PNI
10x
40x
CD45 B220 F4/80 DAPI
e.
Sham
10x
40x
f.
PNI
10x
40x
g.
Macrophages
0.0027
F4/80+ mean gray intensity (a.u)
0
10
20
30
40
50
Sham
PNI
CD45 B220 CD3 DAPI
h.
Sham
10x
40x
i.
PNI
10x
40x

**Figure S5. Immunostaining of B cells with neutrophils, dendritic cells, macrophages, and T cells within $CD45^+$ immune clusters in the DRG meninges.**

Representative 10x and 40x (inset) images of whole mount ipsilateral L4-L5 DRGs collected from male mice 14 days after PNI or sham surgery (scale bar = 100 μm). The enlarged images were digitally cropped from the original 40x images (some are rotated, scale bar = 50 μm). (**a-b**) DRGs immunostained for B cells ($B220^+$), neutrophils ($Ly6G^+$), $CD45^+$ immune cells and DAPI. (**c-d**) DRGs immunostained for dendritic cells ($CD11c^+$), B cells ($B220^+$), $CD45^+$ immune cells and DAPI. (e-**f**) DRGs immunostained for macrophages ($F4/80^+$), B cells ($B220^+$), $CD45^+$ immune cells and DAPI. (**g**) Fluorescence intensity for $F4/80^+$ macrophages were quantified and analyzed by unpaired, two-tailed Student's t test. N = 5-6 per group. **(h-i)** DRGs immunostained for B cells ($B220^+$), T cells ($CD3^+$), $CD45^+$ immune cells and DAPI.

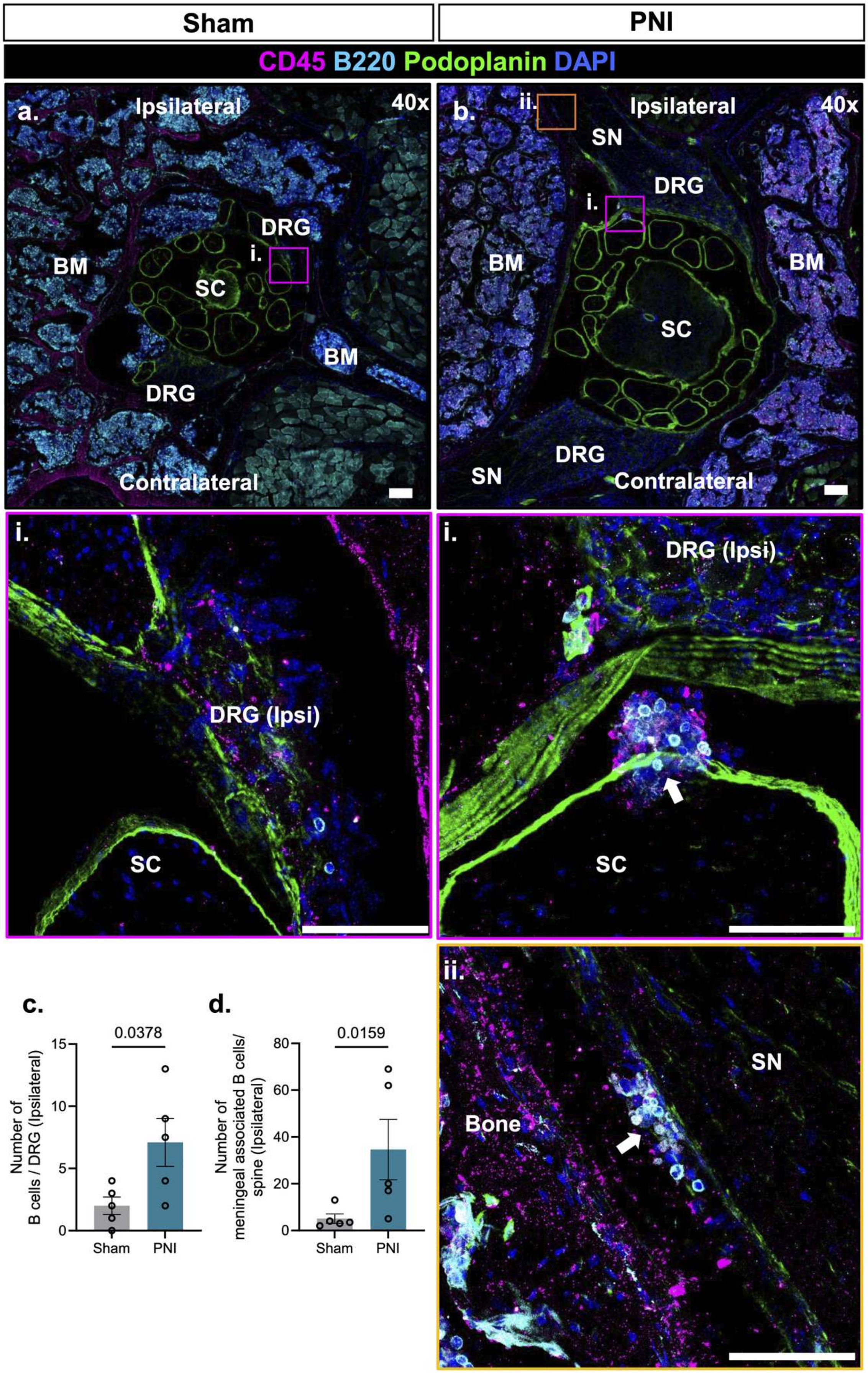

Sham
PNI
CD45 B220 Podoplanin DAPI
a.
Ipsilateral
40x
BM
DRG
i.
SC
BM
DRG
Contralateral
b.
ii.
Ipsilateral
40x
SN
DRG
i.
BM
BM
SC
SN
DRG
Contralateral
i.
DRG (Ipsi)
SC
i.
DRG (Ipsi)
SC
ii.
SN
Bone
c.
0.0378
Number of B cells / DRG (Ipsilateral)
15
10
5
0
Sham
PNI
d.
0.0159
Number of meningeal associated B cells/ spine (Ipsilateral)
80
60
40
20
0
Sham
PNI

**Figure S6. PNI facilitates migration of B cells from the lumbar vertebral bone marrow for TLS development.**

Representative 40x tile images of the L4-L5 region of the spinal column collected from male mice 14 days after PNI. Spine sections were immunostained for B cells (B220), pan-leukocytes (CD45), lymphatic endothelial cells (podoplanin), and DAPI (nuclei) (scale bar = 100 µm). The enlarged images were digitally cropped from the original 40x images (scale bar = 50 µm). (**a, i**) Representative images from sham spinal sections stained for were immunostained for B cells (B220), pan-leukocytes (CD45), lymphatic endothelial cells (podoplanin), and DAPI (nuclei). **(b, i)** Formation of $CD45^{+}B220^{+}$ immune cell cluster at the meningeal interface of the DRG and spinal cord at the ipsilateral side. (**b, ii**) $CD45^{+}B220^{+}$ immune cell cluster associated with the ipsilateral sciatic nerve, adjacent to the DRG. BM: bone marrow; SC: spinal cord, SN; sciatic nerve. (**c-d**) Quantitative analysis of the number of $B220^{+}$ B cells in the ipsilateral side of the spinal DRG and the number of meningeal associated B cells in the ipsilateral side of the spinal sections. N = 5 per group. Data was analyzed using a two-tailed Student's t-test.

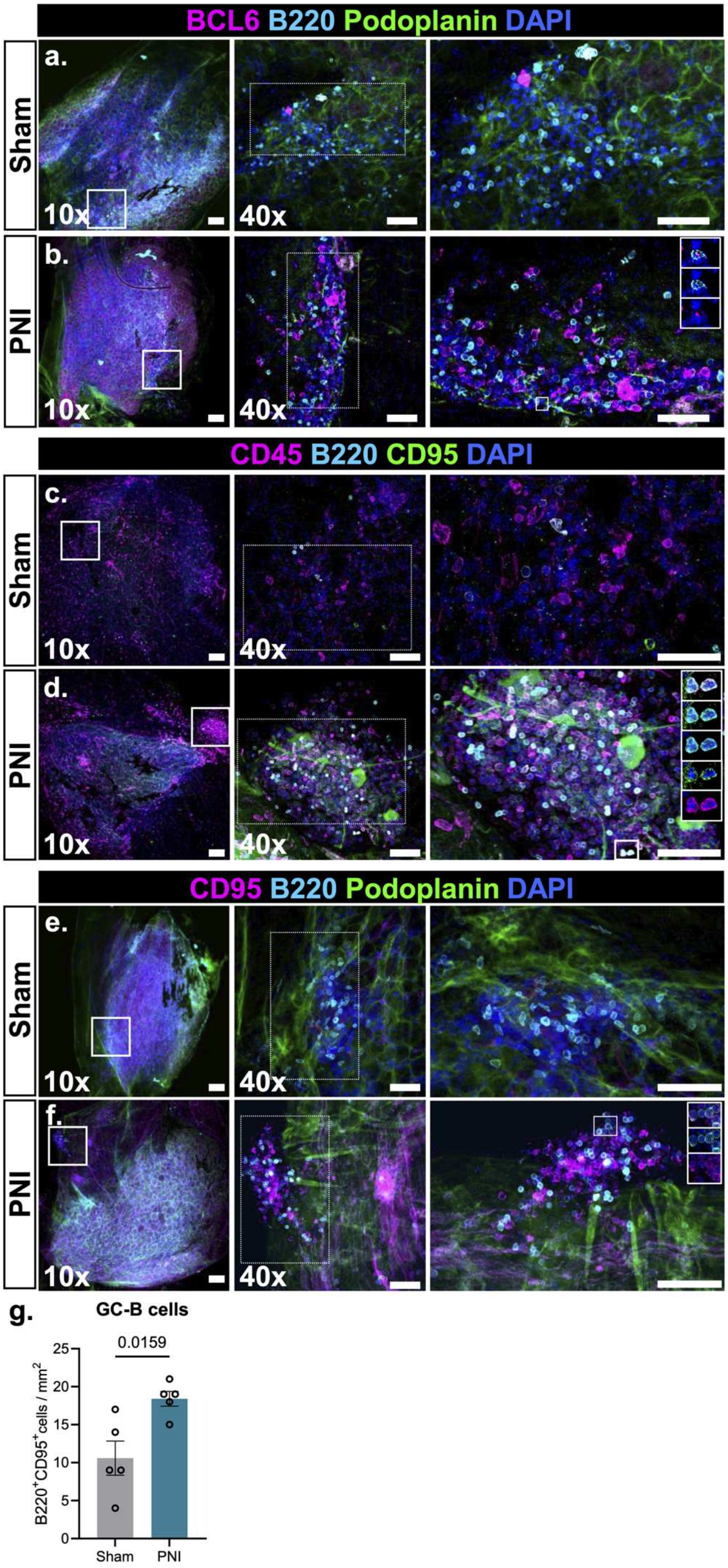

BCL6 B220 Podoplanin DAPI
a.
Sham
10x
40x
b.
PNI
10x
40x
CD45 B220 CD95 DAPI
c.
Sham
10x
40x
d.
PNI
10x
40x
CD95 B220 Podoplanin DAPI
e.
Sham
10x
40x
f.
PNI
10x
40x
g.
GC-B cells
0.0159
B220+CD95+cells / mm2
25
20
15
10
5
0
Sham
PNI

**Figure S7. PNI induces germinal center like structure in the DRG meninges.**
Representative 10x and 40x (inset) images of whole mount contralateral L4-L5 DRGs collected from male mice 14 days after PNI or sham surgery (scale bar = 100 μm). The enlarged images were digitally cropped from the original 40x images (some are rotated, scale bar = 50 μm). (**a-b**) DRGs were stained for B cell ($B220^+$), the germinal center marker $BCL6^+$, lymphatic endothelial marker ($podoplanin^+$) and a nuclear marker (DAPI) to identify germinal center B cells. (**c-d**) DRGs were stained for B cell ($B220^+$), the germinal center marker $CD95^+$, pan leukocyte marker $CD45^+$ and a nuclear marker (DAPI) to identify germinal center B cells. (**e-f**) DRGs were stained for B cell ($B220^+$), the germinal center marker $CD95^+$, lymphatic endothelial marker ($podoplanin^+$) and a nuclear marker (DAPI) to identify germinal center B cells. (**g**) Quantification of $B220^+CD95^+$ germinal center B cells in sham and PNI groups. Data was analyzed using a two-tailed Student's t-test (N = 5 per group).

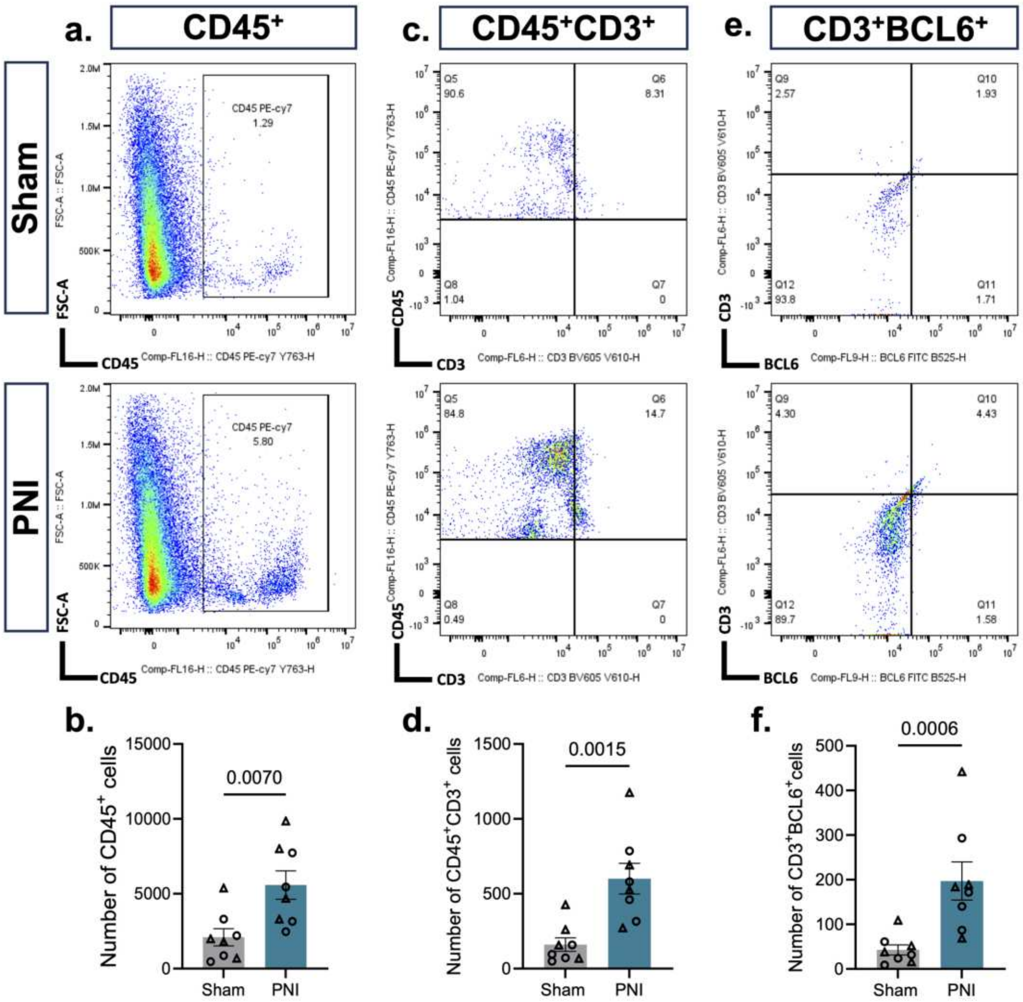


**Figure S8. Flow cytometric analysis of CD45$^{+}$ immune cells, T cells and Tfh cells**.

(**a-b**) Flow cytometric analysis of CD45$^{+}$ immune cells, (**c-d**) CD45$^{+}$CD3$^{+}$ T cells, and (**e-f**) CD45$^{+}$CD3$^{+}$BCL6$^{+}$ Tfh cells. N = 8 samples in each group (DRGs from total 12 male and 12 female mice in each group). Data was analyzed using a two-tailed Student's t-test.

**a.**

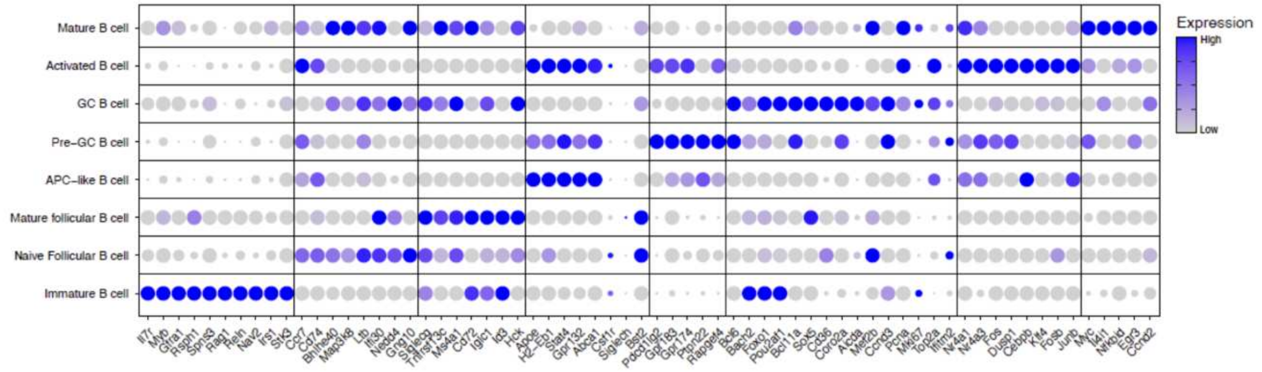


**Figure S9. B cell clustering.**

(**a**) Expression of gene transcripts used to identify B cell subsets from single-cell RNA sequencing of CD45+ cells in the dorsal root ganglia meninges of naïve mice and after PNI (spared nerve injury, second model of PNI).

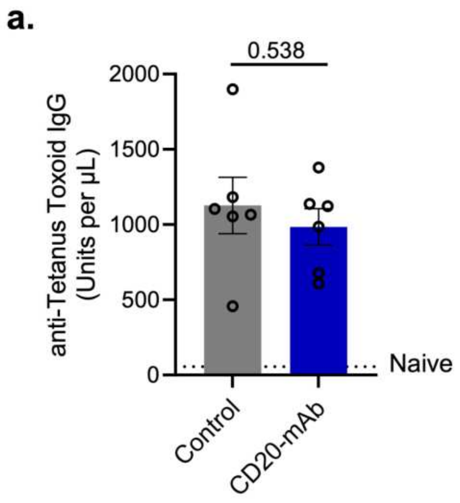


**Figure S10. Serum anti-tetanus IgG levels.**
(**a**) Primary tetanus toxoid vaccine was administered 14 days prior to PNI surgery/CD20 mAb (or isotype control) injections. Booster injections were administered 7 days after PNI surgery/CD20 mAb injections, with secondary IgG responses evaluated 14 days later. Serum anti-tetanus IgG levels were quantified by ELISA. Naïve mice were used as negative control, with titers indicated by the dotted line. Data analyzed by unpaired, two-tailed Student’s t test. N = 3 male and 3 female mice per group.

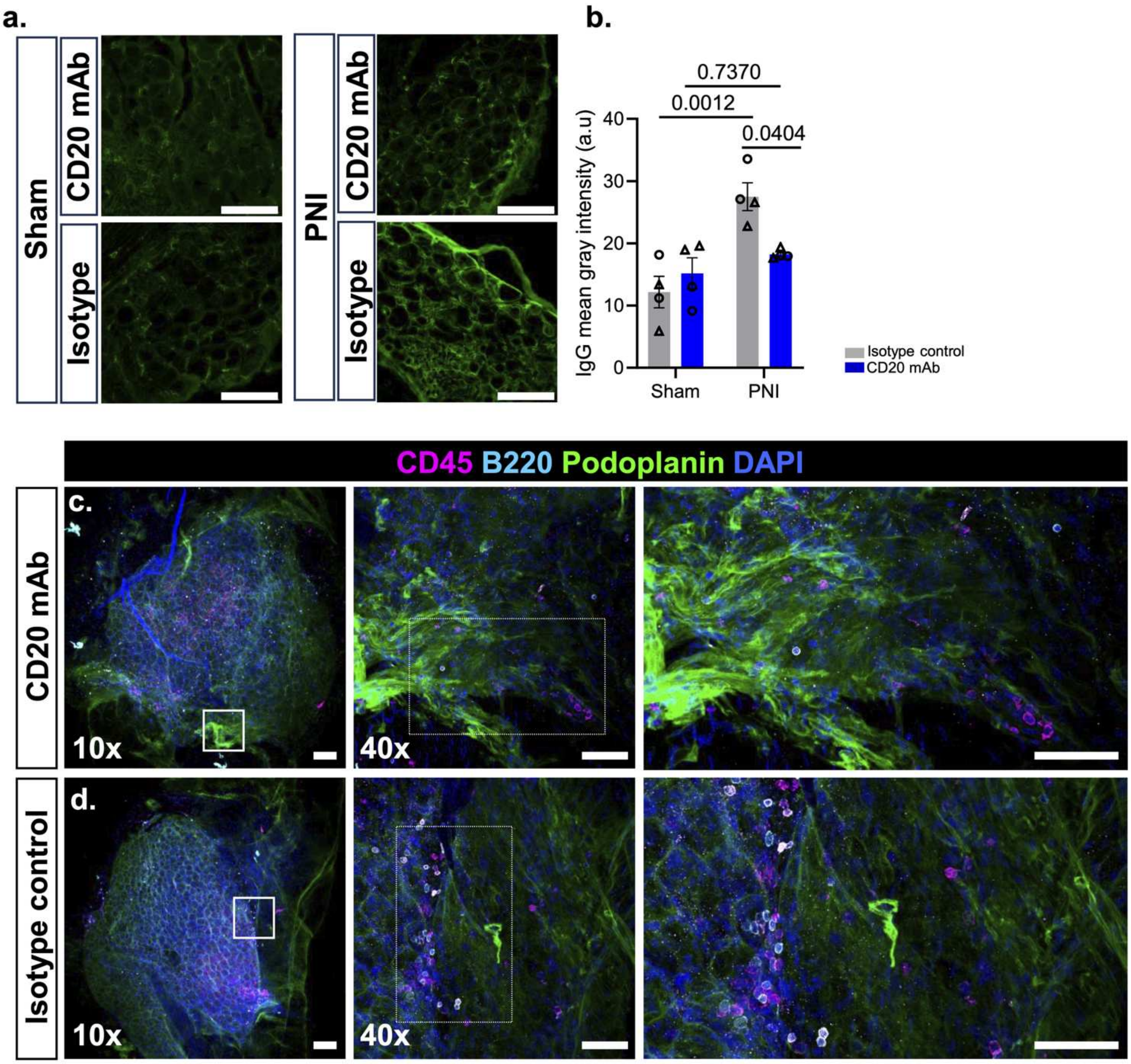


**Figure S11. CD20 mAb reduces IgG deposition in the DRG parenchyma and meningeal TLSs do not develop in sham DRGs and are not altered by CD20 mAb treatment.**

(**a**) Representative 40x images of DRG sections stained for IgG from sham control treated with CD20 mAb or isotype control antibody, along with PNI mice. (scale bar = 100 µm) **(b)** Quantification of IgG mean gray intensity in DRG sections across groups. Data was analyzed by two-way ANOVA and Tukey's multiple-comparisons test (N = 4 mice per group; both sexes included). (**c-d**) Representative 10x and 40x (inset) images of whole mount ipsilateral L4-L5 DRGs collected 28 days after sham surgery and intrathecal CD20 mAb or isotype control antibody treatment (scale bar = 100 µm). The enlarged images were digitally cropped from the original 40x images (some are rotated, scale bar = 50 µm). DRGs were immunostained for B cells (B220), pan-leukocytes (CD45), lymphatic endothelial cells (podoplanin), and DAPI (nuclei).

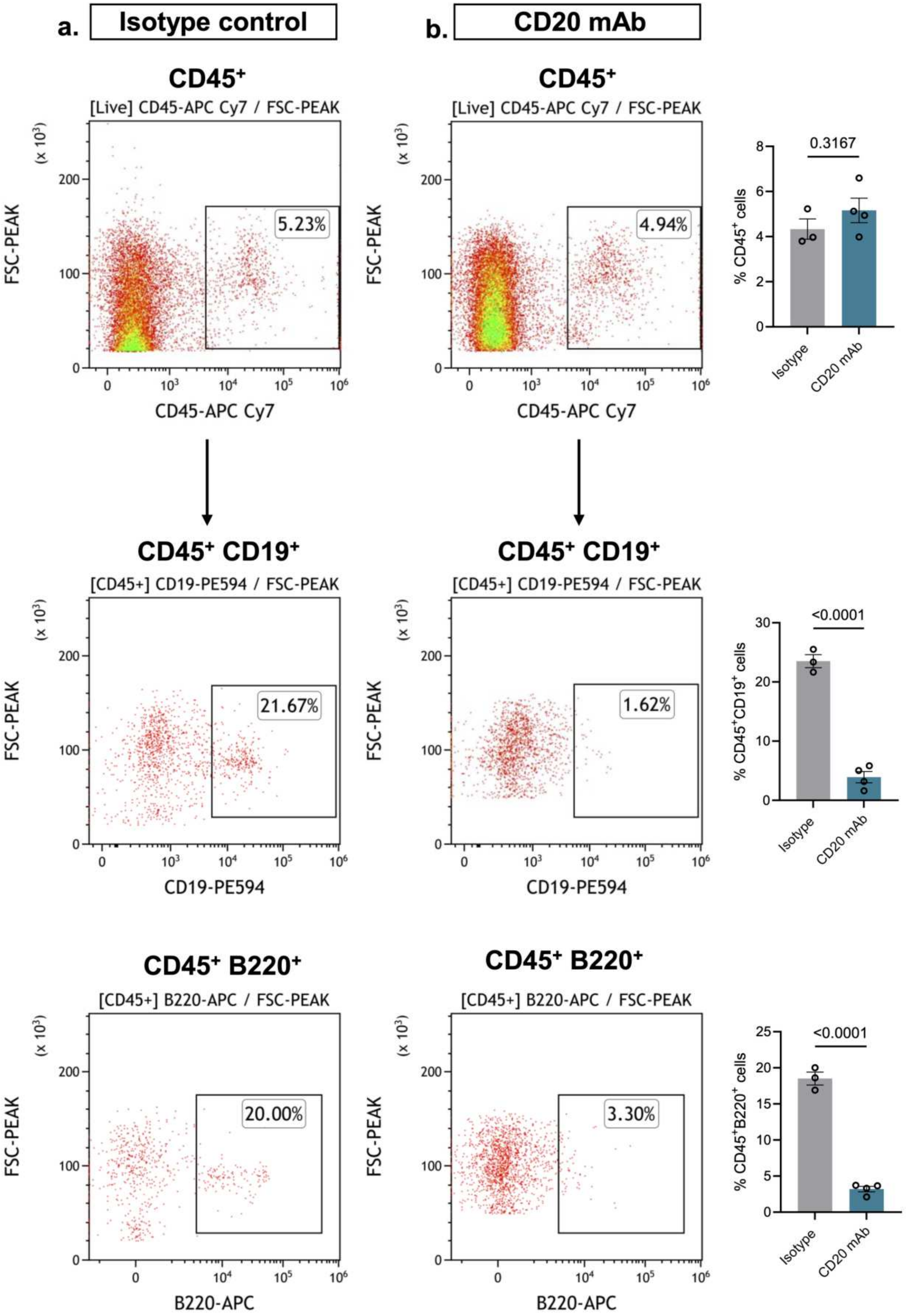


**Figure S12. Flow cytometric analysis of B cells in the DRGs following CD20 mAb treatment.**

(**a–b**) Flow cytometric analysis of cells isolated from DRGs with meninges 14 days after PNI from mice treated with CD20 mAb or isotype control antibody. Representative flow cytometry plots showing $CD45^+$ cells, $CD45^+CD19^+$ cells, and $CD45^+B220^+$ cells from DRGs of (a) PNI + isotype control and (b) PNI + CD20 mAb-treated mice. Quantification of the percentage of $CD45^+$, $CD45^+CD19^+$, and $CD45^+B220^+$ cells is shown. Data were analyzed using a two-tailed Student's t-test (N = 3–4 samples per group, DRGs from total 6-8 mice in each group).

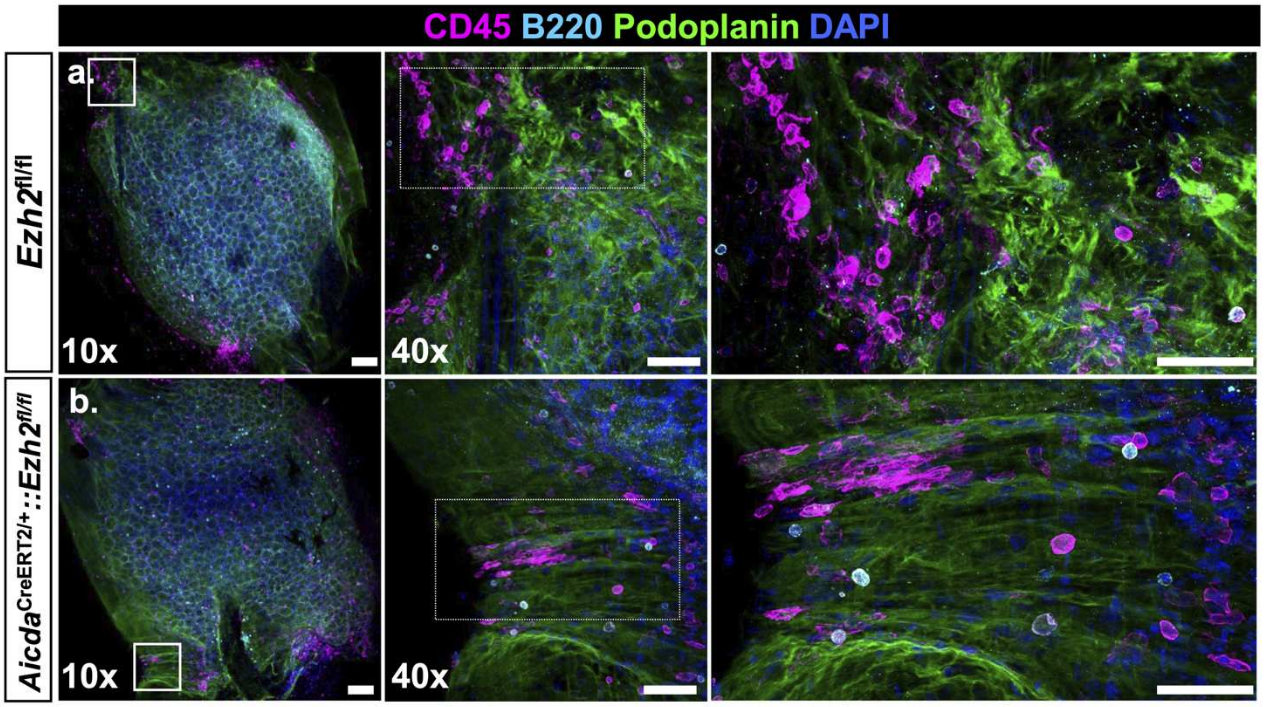


**Figure S13**. **Meningeal TLSs do not develop in sham DRGs of *Aicda*$^{creERT2/+}$::*Ezh2*$^{fl/fl}$ mice.**
**(a-b)** Representative 10x and 40x (inset) images of whole mount ipsilateral L4-L5 DRGs collected 28 days after sham surgery in *Aicda*$^{creERT2/+}$::*Ezh2*$^{fl/fl}$ mice or littermate control mice (scale bar = 100 μm). The enlarged images were digitally cropped from the original 40x images (scale bar = 50 μm**)**. DRGs were immunostained for B cells (B220), pan-leukocytes (CD45), lymphatic endothelial cells (podoplanin), and DAPI (nuclei).

a.

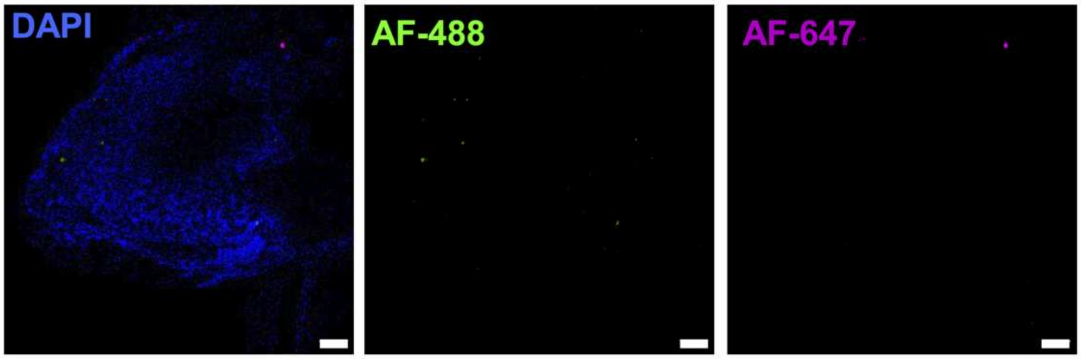


**Figure S14. Controls for immunostaining in whole mount DRG.**
(**a**) Representative images acquired from DRG images immunostained for the secondary antibodies alone (Alexa Fluor 488 and Alexa Fluor 647). Scale bar = 100 µm.

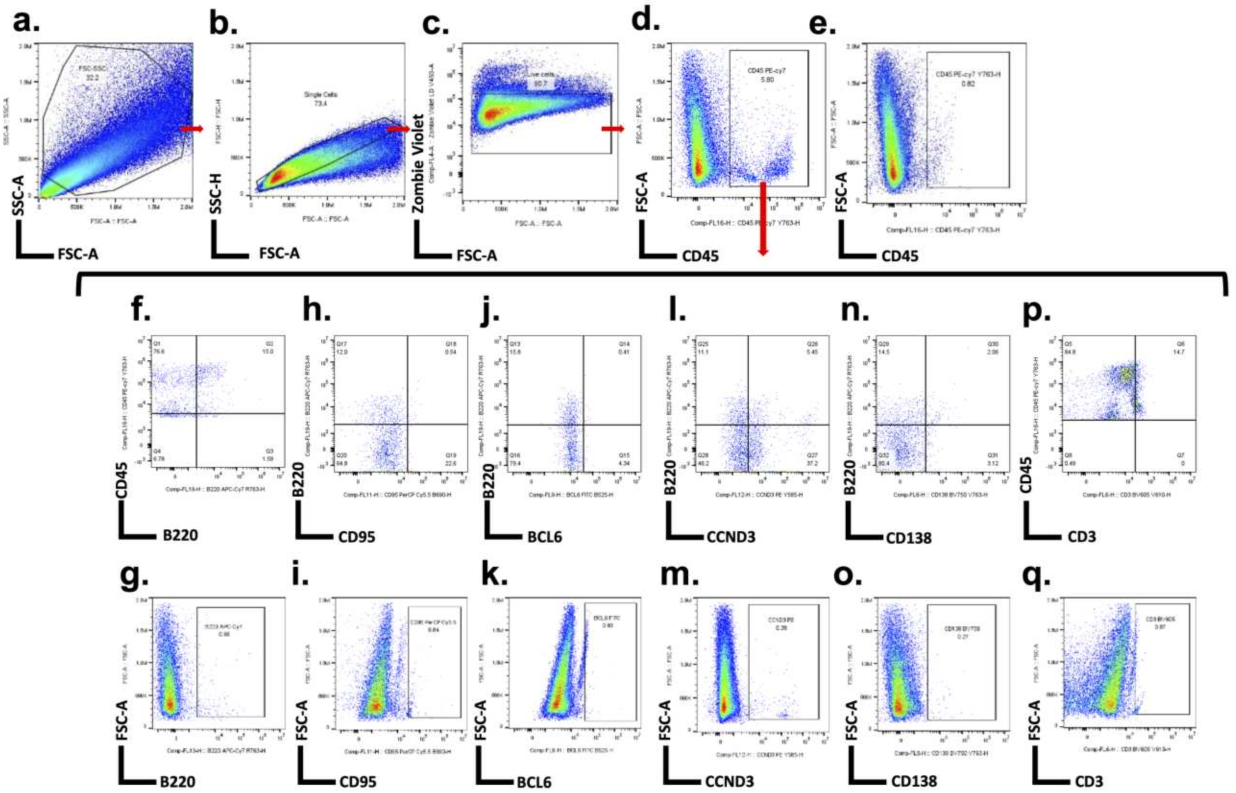


**Figure S15. Representative gating strategy for flow cytometric analysis of germinal center associated markers.**

Representative flow cytometry plots illustrating the sequential gating strategy used for analysis of DRG meningeal immune cell populations and germinal center–associated markers. Cells were first gated based on **(a)** forward scatter (FSC) versus side scatter (SSC) to exclude debris, followed by **(b)** singlet discrimination to remove aggregates and **(c)** live-cell gating to exclude dead cells. From the live-cell population, **(d–e)** $CD45^+$ leukocytes were identified with the corresponding fluorescence minus one (FMO) control. Within the $CD45^+$ population, the following subsets were further analyzed with their respective FMO controls: **(f–g)** $B220^+$ B cells and $B220^+$ FMO; **(h–i)** $B220^+CD95^+$ cells and $CD95^+$ FMO; **(j-k)** $B220^+BCL6^+$ cells and $BCL6^+$ FMO; **(l-m)** $B220^+CCND3^+$ cells and $CCND3^+$ FMO; **(n-o)** $B220^+CD138^+$ cells and $CD138^+$ FMO; and **(p-q)** $CD45^+CD3^+$ T cells and $CD3^+$ FMO.

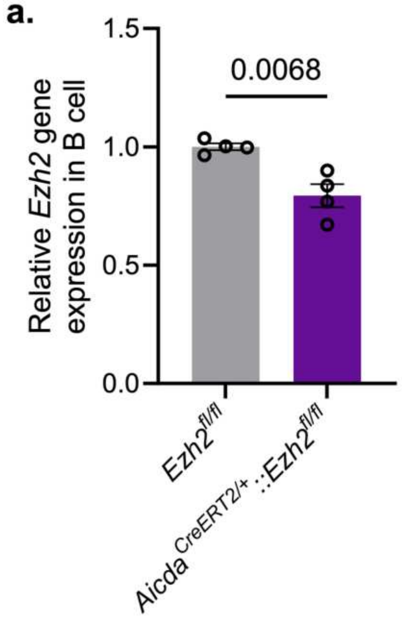


**Figure S16. Validation of *Ezh2* deletion from B cells.**
(**a**) Ezh2 expression in splenic B cells purified from *Aicda*$^{creERT2/+}$::*Ezh2*$^{fl/fl}$ and littermate control mice at 14 days after PNI. Mice received tamoxifen (10 mg/kg, oral gavage) for four consecutive days. Data analyzed by unpaired, two-tailed Student's t test. N = 4 mice per group.

### SUPPLEMENTARY TABLES

**Table S1. List of antibodies**

| Antibodies | Company | Catalogue number | Dilution factor |
|---|---|---|---|
| CD45 | Abcam | AB10558 | 1:250 |
| CD45 | BD Pharmingen | 550539 | 1:100 |
| B220 | Invitrogen | 14-0452-85 | 1:300 |
| CD20 | Abcam | AB64088 | 1:250 |
| CD3 | Invitrogen | 14-0032-82 | 1:300 |
| F4/80 | Cell Signaling Technology | 71299 | 1:250 |
| Ly6G | Abcam | AB25377 | 1:300 |
| PCNA | Novus Biologicals | NB100-456 | 1:200 |
| MECA-79 | Invitrogen | 53-6036-82 | 1:200 |
| BCL6 | Abcam | ab272859 | 1:250 |
| CD11c | Cell Signaling Technology | 97585 | 1:250 |
| Podoplanin | Invitrogen | 14-5381-82 | 1:350 |
| Lyve-1 | Invitrogen | 14-0443-82 | 1:350 |
| GLUT-1 | Abcam | Ab209449 | 1:100 |
| Hamster Alexa Fluor 488 | Invitrogen | A21110 | 1:400 |
| Hamster Alexa Fluor 647 | Invitrogen | A21451 | 1:400 |
| Rat Alexa Fluor 594 | Invitrogen | A11007 | 1:300 |
| Rat Alexa Fluor 488 | Invitrogen | A11006 | 1:300 |
| Rabbit Alexa Fluor 488 | Invitrogen | A11008 | 1:300 |
| Rabbit Alexa Fluor 647 | Invitrogen | A21245 | 1:300 |
| Rabbit Alexa Fluor 594 | Invitrogen | A11012 | 1:300 |
| DAPI | Invitrogen | 62248 | 1:1000 |
| CD21/35 | Abcam | AB318999 | 1:200 |
| PD-1 | Abcam | AB214421 | 1:100 |
| Gr-1 | R&D Systems | MAB1037 | 3μg/ml |
| CD79a (pig) | NeoBiotechnologies | 973-RBM10-P0 | 1:50 |
| IgG (pig) | Jackson ImmunoResearch | 114-005-003 | 1:100 |
| BA4D5 (pig) | Invitrogen | MA5-28472 | 1:50 |
| CD95 | Abcam | AB21088 | 1:100 |
| CD138 | Invitrogen | 36-2900 | 1:300 |
| APC B220 | Biolegend | 103212 | 1:100 |
| APC/Cyanine7 CD45 | Biolegend | 103116 | 1:100 |
| PE/Dazzle™ 594 CD19 | Biolegend | 115554 | 1:100 |
| LIVE/DEAD™ Yellow | Biolegend | 423104 | 1:1000 |
| PE-Cy7 CD45 | Biolegend | 103114 | 1:100 |
| APC-Cy7 CD45R (B220) | Biolegend | 103224 | 1:100 |
| Percp-Cy5.5 CD95 | Biolegend | 152610 | 1:100 |
| FITC BCL6 | Biolegend | 358513 | 1:100 |

| | | | |
|---|---|---|---|
| Alexa Fluor 647 BCL2 | Biolegend | 658705 | 1:100 |
| PE CCND3 | Biolegend | 684904 | 1:100 |
| Brilliant Violet 750 CD138 | BD Biosciences | 747070 | 1:100 |
| Brilliant Violet 605 CD3 | Biolegend | 100237 | 1:100 |
| Zombie Violet | Biolegend | 77477 | 1:1000 |

**Table S2. Morphometric parameters of CD45$^+$ immune cell clusters**

| | **Small aggregates or single cells** | **Medium clusters** | **Large clusters** |
|---|---|---|---|
| Surface Area (μm$^2$); median (range) | 464 (300 – 14644) | 4701 (1376 – 48884) | 12477 (2363 – 156030) |
| Volume (μm$^3$); median (range) | 396 (56 – 41981) | 6089 (1105 – 92755) | 14644 (2081 – 88841) |
| Sphericity; median (range) | 0.53 (0.02 – 0.97) | 0.33 (0.04 – 0.59) | 0.25 (0.06 – 0.56) |
| Ellipticity (oblate); median (range) | 0.35 (0.01 – 0.94) | 0.38 (0.00 – 0.89) | 0.39 (0.08 – 0.86) |
| Ellipticity (prolate); median (range) | 0.31 (0.00 – 0.94) | 0.14 (0.00 – 0.70) | 0.06 (0.00 – 0.32) |

**Table S3: Gene list for TLS analysis in human DRGs**

| **hgnc_symbol** | **gene** | **gs** | **ensembl_gene_id** |
|---|---|---|---|
| AICDA | AICDA | TLS_curated gene list | ENSG00000111732 |
| BCL6 | BCL6 | TLS_curated gene list | ENSG00000113916 |
| CD19 | CD19 | TLS_curated gene list | ENSG00000177455 |
| CD3E | CD3E | TLS_curated gene list | ENSG00000198851 |
| CD4 | CD4 | TLS_curated gene list | ENSG00000010610 |
| CD79A | CD79A | TLS_curated gene list | ENSG00000105369 |
| CXCL13 | CXCL13 | TLS_curated gene list | ENSG00000156234 |
| CXCR5 | CXCR5 | TLS_curated gene list | ENSG00000160683 |
| FAS | FAS | TLS_curated gene list | ENSG00000026103 |
| ICOS | ICOS | TLS_curated gene list | ENSG00000163600 |

| | | | |
|---|---|---|---|
| LTBR | LTBR | TLS_curated gene list | ENSG00000111321 |
| MEF2B | MEF2B | TLS_curated gene list | ENSG00000213999 |
| MKI67 | MKI67 | TLS_curated gene list | ENSG00000148773 |
| PCNA | PCNA | TLS_curated gene list | ENSG00000132646 |
| PDCD1 | PDCD1 | TLS_curated gene list | ENSG00000276977 |
| PDCD1 | PDCD1 | TLS_curated gene list | ENSG00000188389 |
| PRDM1 | PRDM1 | TLS_curated gene list | ENSG00000057657 |
| TOP2A | TOP2A | TLS_curated gene list | ENSG00000131747 |
| APOE | APOE | TLS_Meylan2022 | ENSG00000130203 |
| C1QA | C1QA | TLS_Meylan2022 | ENSG00000173372 |
| C7 | C7 | TLS_Meylan2022 | ENSG00000112936 |
| CD52 | CD52 | TLS_Meylan2022 | ENSG00000169442 |
| CD79A | CD79A | TLS_Meylan2022 | ENSG00000105369 |
| CXCL12 | CXCL12 | TLS_Meylan2022 | ENSG00000107562 |
| DERL3 | DERL3 | TLS_Meylan2022 | ENSG00000099958 |
| DERL3 | DERL3 | TLS_Meylan2022 | ENSG00000274437 |
| FCRL5 | FCRL5 | TLS_Meylan2022 | ENSG00000143297 |
| IGHA1 | IGHA1 | TLS_Meylan2022 | ENSG00000211895 |
| IGHA1 | IGHA1 | TLS_Meylan2022 | ENSG00000282633 |
| IGHG1 | IGHG1 | TLS_Meylan2022 | ENSG00000211896 |
| IGHG1 | IGHG1 | TLS_Meylan2022 | ENSG00000277633 |
| IGHG2 | IGHG2 | TLS_Meylan2022 | ENSG00000274497 |
| IGHG2 | IGHG2 | TLS_Meylan2022 | ENSG00000211893 |
| IGHG3 | IGHG3 | TLS_Meylan2022 | ENSG00000211897 |
| IGHG3 | IGHG3 | TLS_Meylan2022 | ENSG00000282184 |
| IGHG4 | IGHG4 | TLS_Meylan2022 | ENSG00000277016 |
| IGHG4 | IGHG4 | TLS_Meylan2022 | ENSG00000211892 |
| IGHGP | IGHGP | TLS_Meylan2022 | ENSG00000253755 |
| IGHGP | IGHGP | TLS_Meylan2022 | ENSG00000282094 |
| IGHM | IGHM | TLS_Meylan2022 | ENSG00000282657 |
| IGHM | IGHM | TLS_Meylan2022 | ENSG00000211899 |
| IGKC | IGKC | TLS_Meylan2022 | ENSG00000211592 |
| IGLC1 | IGLC1 | TLS_Meylan2022 | ENSG00000211675 |
| IGLC2 | IGLC2 | TLS_Meylan2022 | ENSG00000211677 |
| IGLC3 | IGLC3 | TLS_Meylan2022 | ENSG00000211679 |

| | | | |
|---|---|---|---|
| IL7R | IL7R | TLS_Meylan2022 | ENSG00000168685 |
| JCHAIN | JCHAIN | TLS_Meylan2022 | ENSG00000132465 |
| LUM | LUM | TLS_Meylan2022 | ENSG00000139329 |
| MZB1 | MZB1 | TLS_Meylan2022 | ENSG00000170476 |
| PIM2 | PIM2 | TLS_Meylan2022 | ENSG00000292210 |
| PIM2 | PIM2 | TLS_Meylan2022 | ENSG00000102096 |
| PTGDS | PTGDS | TLS_Meylan2022 | ENSG00000107317 |
| SSR4 | SSR4 | TLS_Meylan2022 | ENSG00000180879 |
| TRBC2 | TRBC2 | TLS_Meylan2022 | ENSG00000211772 |
| TRBC2 | TRBC2 | TLS_Meylan2022 | ENSG00000276849 |
| XBP1 | XBP1 | TLS_Meylan2022 | ENSG00000100219 |
